# Novel torin1-sensitive phosphorylation sites on the metabolic regulator AMPK revealed by label-free mass spectrometry

**DOI:** 10.1101/2021.10.20.462995

**Authors:** William J. Smiles, Ashley J. Ovens, Dingyi Yu, Naomi X.Y. Ling, Kaitlin R. Morrison, Ashfaqul Hoque, John W. Scott, Sandra Galic, Christopher G. Langendorf, Bruce E. Kemp, Janni Peterson, Jonathan S. Oakhill

**Affiliations:** Metabolic Signalling Laboratory, St. Vincent’s Institute of Medical Research, Fitzroy 3065, Australia; Mary MacKillop Institute for Health Research, Australian Catholic University, Melbourne 3000, Australia; Protein Chemistry and Metabolism, St. Vincent’s Institute of Medical Research, Fitzroy 3065, Australia; Flinders Health and Medical Research Institute, Flinders Centre for Innovation in Cancer, Flinders University, Adelaide, South Australia, Australia; The Florey Institute of Neuroscience and Mental Health, Royal Parade, Parkville 3052, Australia

## Abstract

AMPK and mTORC1 are nutrient-sensitive protein kinases that form a fundamental negative feedback loop that governs cell growth and proliferation. AMPK is an αβγ heterotrimer that is directly phosphorylated by mTORC1 on α2^S345^ to suppress AMPK activity and promote cell proliferation under nutrient stress conditions. Using mass spectrometry, we generated precise phosphorylation profiles of all 12 AMPK complexes expressed in proliferating human cells. Of the 18 phosphorylation sites detected, seven were sensitive to pharmacological mTORC1 inhibition, including four in the AMPK γ2 isoform NH_2_-terminal domain and α2^S377^ which is located in the nucleotide-sensing motif. In particular, β1^S182^ and β2^S184^ were found to be mTORC1 substrates *in vitro* and near-maximally or substantially phosphorylated under cellular growth conditions. β^S182^ phosphorylation was elevated in α1-containing complexes, relative to α2, an effect partly attributable to the non-conserved α-subunit serine/threonine-rich loop. While mutation of β1^S182^ to a non-phosphorylatable Ala had no effect on basal and ligand-stimulated AMPK activity, β2-S184A mutation increased nuclear AMPK activity and enhanced cell proliferation under nutrient stress. We conclude that mTORC1 governs the nuclear activity of AMPK to regulate transcription factors involved in metabolism and cell survival during nutrient shortage.

## Introduction

Central to the regulation of cellular metabolism are signalling networks propagated by diverse molecular pathways that couple the direct sensing of metabolites to enzyme catalysis and the covalent modification of proteins with either small (e.g., phosphate groups) or large (e.g., ubiquitin) attachments [1–3]. Fundamental to this regulation of metabolism are the nutrient-sensitive Ser/Thr protein kinases AMP-activated protein kinase (AMPK) and mechanistic (formerly ‘mammalian’) target of rapamycin complex 1 (mTORC1), which, broadly speaking, antagonistically drive catabolic (e.g., autophagy, lipolysis) and anabolic (e.g., protein and ribosome synthesis) processes, respectively. AMPK is considered the energy guardian of the cell due to its role in maintaining cellular energy balance [4,5], forming heterotrimeric complexes comprising an α catalytic subunit, as well as β and γ regulatory subunits [6]. Multiple isoforms of each subunit exist (α1/2, β1/2, γ1/2/3), allowing for up to 12 unique AMPK complexes to form, each with distinct tissue distribution patterns [6,7].

The α-subunit kinase domain (α-KD) possesses an archetypal kinase domain structure with an activation loop phosphorylation site α1^T174^/α2^T172^ (α^T172^) targeted by liver kinase B1 (LKB1) and calcium/calmodulin-dependent kinase kinase 2 (CaMKK2) [8–11]. Additional COOH-terminal regulatory elements on the α-subunit include a tri-helical autoinhibitory domain (AID), α-regulatory subunit-interacting motifs (α-RIM) that interact with the γ-subunit, and a serine/threonine-rich loop (ST loop) heavily modified by phosphorylation [12]. The β subunit is capable of anchoring to carbohydrates through its carbohydrate binding module (CBM), and it contains a COOH-terminal α-γ-subunit binding sequence critical for complex formation [13]. In the quaternary AMPK structure, the CBM docks onto the α-KD N-lobe to form a hydrophobic cleft termed the allosteric drug and metabolite site (ADaM site) allowing allosteric activation via synthetic compounds (e.g., SC4, MSG011, MK8722, PF739) and endogenous long chain fatty acyl-CoA esters (e.g., palmitoyl-CoA) [14–17]. However, the most well-known mechanism of AMPK activation is through elevations in AMP/ATP and ADP/ATP ratios, where adenine nucleotides directly bind to the γ-subunit at sites formed by four cystathionine β-synthase repeats [13]. The presence of nucleotides bound at the γ subunit is sensed by the α-RIM, where binding of AMP sequesters the AID from the α-KD to allosterically activate AMPK, whereas AMP or ADP binding also promote α^T172^ phosphorylation (p-α^T172^) and protect it from dephosphorylation [18,19]. However, p-α^T172^ is not absolutely required for AMPK activity *in vitro* and *in cellulo*, where significant synergist allosteric activation of up to 1000-fold can be achieved with dephosphorylated α^T172^ via dual ligand binding of an ADaM site compound with AMP [20]. Given AMPK’s central role in regulating metabolism, it is unsurprising that it is also subject to multidirectional regulatory inputs via a range of phosphorylation sites present across each of its subunits [12].

mTOR is the catalytic component of mTORC1 and mTORC2, two structurally and functionally distinct multi-protein complexes defined by unique regulatory partners (mTORC1, Raptor; mTORC2, Rictor/SIN1) that dictate substrate selectivity. mTORC1 is activated by growth factors and nutrients such as amino acids, which canonically harness the lysosome as the cellular locale integrating these physiological cues to stimulate mTORC1 [21–24]. AMPK inhibits mTORC1 directly and indirectly by phosphorylation of Raptor and TSC2, respectively [25,26]. Recently, we found that mTORC1 directly inhibits AMPK activity in yeast and mammalian cells by phosphorylating α1^S347^/α2^S345^ (p-α^S345^), however the precise mechanism remains unknown [27].

Examinations of cellular signalling have traditionally relied heavily on semi-quantitative immunoblotting techniques that are often fraught with technical limitations (e.g., phospho-antibody linearity/sensitivity). Here, we circumvent these issues by taking advantage of a highly targeted mass spectrometry (MS)-based approach to determine the basal phosphorylation stoichiometries of 18 phosphorylation sites (10 novel and 13 with unknown function) among all 12 AMPK complexes expressed in mammalian cells. Notably, β1^S182^/β2^S184^ (β^S182^) is the most heavily phosphorylated site on AMPK, and along with six other phosphorylation sites, four of which are located in the γ2-unique NH_2_-terminal extension (NTE), is sensitive to mTOR inhibition *in cellulo*. We confirmed β^S182^ as a direct mTORC1 substrate, phosphorylation of which inhibits nuclear AMPK activity of complexes containing the β2 isoform, in turn restricting cell proliferation under nutrient stress.

## Results

### Determining phosphorylation profiles across all AMPK heterotrimers

To gain a more comprehensive view of the AMPK phosphoprofile landscape and identify isoform-specific differences, we expressed all 12 AMPK heterotrimers (FLAG-tagged) in a highly proliferative human cell line (HEK293T/17) in complete growth media. FLAG-immunoprecipitated AMPK was then subjected to on-resin tryptic digestion and peptides were analysed by liquid chromatography-mass spectrometry (LC-MS). A targeted approach was employed to achieve optimal sequence coverage of all seven AMPK subunits permitting detection of an assortment of known and novel phosphorylation sites. With precision we could quantify the stoichiometry of these sites (Figure 2 & Table 1) using both phosphorylated and dephosphorylated peptide peak areas (Equation 1). However, our analysis failed to detect some well-known AMPK phosphorylation sites, such as α^T172^ and autophosphorylation residues β1^S108^ and α2^S491^. We speculate the inability to identify p-α^T172^ in our targeted MS approach was due to a salt bridge forming between the α^T172^ phosphate and α^R171^ at the P-1 position that obstructed tryptic cleavage [28,29].

**Table 1.** LC-MS determined stoichiometries of phosphorylated residues identified on AMPK expressed in HEK293T/17 cells. Values represent average stoichiometries from three replicates. SEM values ranged from 0.03 (p-β2^S108^ on α2β2γ1) to 1.69 (p-α2^S345^ on α2β2γ2).

| AMPK complex |  |  |  |  |  |  |  |  |  |  |  |  |
| --- | --- | --- | --- | --- | --- | --- | --- | --- | --- | --- | --- | --- |
| | $\alpha 1\beta 1\gamma 1$ | $\alpha 1\beta 1\gamma 2$ | $\alpha 1\beta 1\gamma 3$ | $\alpha 2\beta 1\gamma 1$ | $\alpha 2\beta 1\gamma 2$ | $\alpha 2\beta 1\gamma 3$ | $\alpha 1\beta 2\gamma 1$ | $\alpha 1\beta 2\gamma 2$ | $\alpha 1\beta 2\gamma 3$ | $\alpha 2\beta 2\gamma 1$ | $\alpha 2\beta 2\gamma 2$ | $\alpha 2\beta 2\gamma 3$ |
| p-$\alpha 1/2^{S347/S345}$ | 41.79 | 44.54 | 39.07 | 66.38 | 59.15 | 69.77 | 33.83 | 45.33 | 35.41 | 63.95 | 58.73 | 59.30 |
| p-$\alpha 1/2^{T373/S377}$ | no phos. detected | no phos. detected | no phos. detected | 22.05 | 18.01 | 27.30 | no phos. detected | no phos. detected | no phos. detected | 15.13 | 16.98 | 23.97 |
| p-$\alpha 1/2^{S477/S481}$ | 7.03 | 9.91 | 5.35 | 3.57 | 12.45 | 4.78 | 6.47 | 9.67 | 4.82 | 6.58 | 18.24 | 7.81 |
| p-$\alpha 1/2^{T481/T485}$ | 22.10 | 29.33 | 22.36 | 8.45 | 14.50 | 13.49 | 22.58 | 31.25 | 21.34 | 11.03 | 19.58 | 17.57 |
| p-$\alpha 1/2^{S487/S491}$ | 32.27 | 32.95 | 20.35 | n.d. | n.d. | n.d. | 28.68 | 33.30 | 18.68 | n.d. | n.d. | n.d. |
| p-$\alpha 1^{S499}$ | 9.28 | 15.60 | 14.30 | - | - | - | 8.46 | 17.16 | 10.83 | - | - | - |
| p-$\alpha 2^{S501}$ | - | - | - | 6.92 | 8.17 | 7.57 | - | - | - | 8.67 | 10.36 | 9.92 |
| p-$\beta^{S108}$ | n.d. | n.d. | n.d. | n.d. | n.d. | n.d. | 5.51 | 29.22 | 5.97 | 3.92 | 26.42 | 19.26 |
| p-$\beta 1/2^{S182/184}$ | 94.07 | 90.68 | 88.09 | 95.38 | 86.01 | 93.27 | 96.43 | 92.51 | 93.30 | 87.16 | 81.15 | 79.91 |
| p-$\gamma 2^{S113}$ | - | 27.59 | - | - | 24.91 | - | - | 24.28 | - | - | 21.17 | - |
| p-$\gamma 2^{S143}$ | - | 15.68 | - | - | 15.71 | - | - | 15.72 | - | - | 15.07 | - |
| p-$\gamma 2^{S162}$ | - | 13.12 | - | - | 13.06 | - | - | 11.13 | - | - | 10.27 | - |
| p-$\gamma 2^{S196}$ | - | 56.57 | - | - | 55.05 | - | - | 50.46 | - | - | 50.29 | - |
| p-$\gamma 2^{133-148}$ Peak 2 | - | 3.01 | - | - | 2.78 | - | - | 2.25 | - | - | 2.31 | - |
| p-$\gamma 2^{156-171}$ Peak 1 | - | 1.48 | - | - | 1.56 | - | - | 1.29 | - | - | 1.66 | - |
| p-$\gamma 2^{156-171}$ Peak 3 | - | 3.77 | - | - | 3.73 | - | - | 2.84 | - | - | 3.09 | - |
| p-$\gamma 3^{S14}$ | - | - | 61.38 | - | - | 61.51 | - | - | 64.10 | - | - | 65.64 |
| p-$\gamma 3^{S65}$ | - | - | 2.47 | - | - | 3.41 | - | - | 2.78 | - | - | 1.99 |

Phosphorylation (%)

These absent phosphosites were detected using validated, site-specific antibodies by immunoblotting of a small volume of FLAG-immunoprecipitated AMPK (Figure 1A). Whilst immunoblotting precluded exact determination of phosphorylation stoichiometries, important isoform-specific differences were still observable. Our immunoblot data suggests that in HEK293T/17 cells, α1^T174^ is more heavily phosphorylated than the equivalent α2 (Figure 1B), which is consistent with AMPK purified from rat liver [8]. Indeed, p-α2^T172^ has been shown to be more sensitive to phosphatases than p-α1^T172^ [30], suggesting greater turnover of the former in mammalian cells. Our results also demonstrate that γ2-AMPK complexes have significantly higher basal p-α^T172^ versus γ1- and γ3-complexes (Figure 1B), in agreement with previous findings showing that the γ2-NTE provides phosphatase protection [19,31].

**Figure 1.**
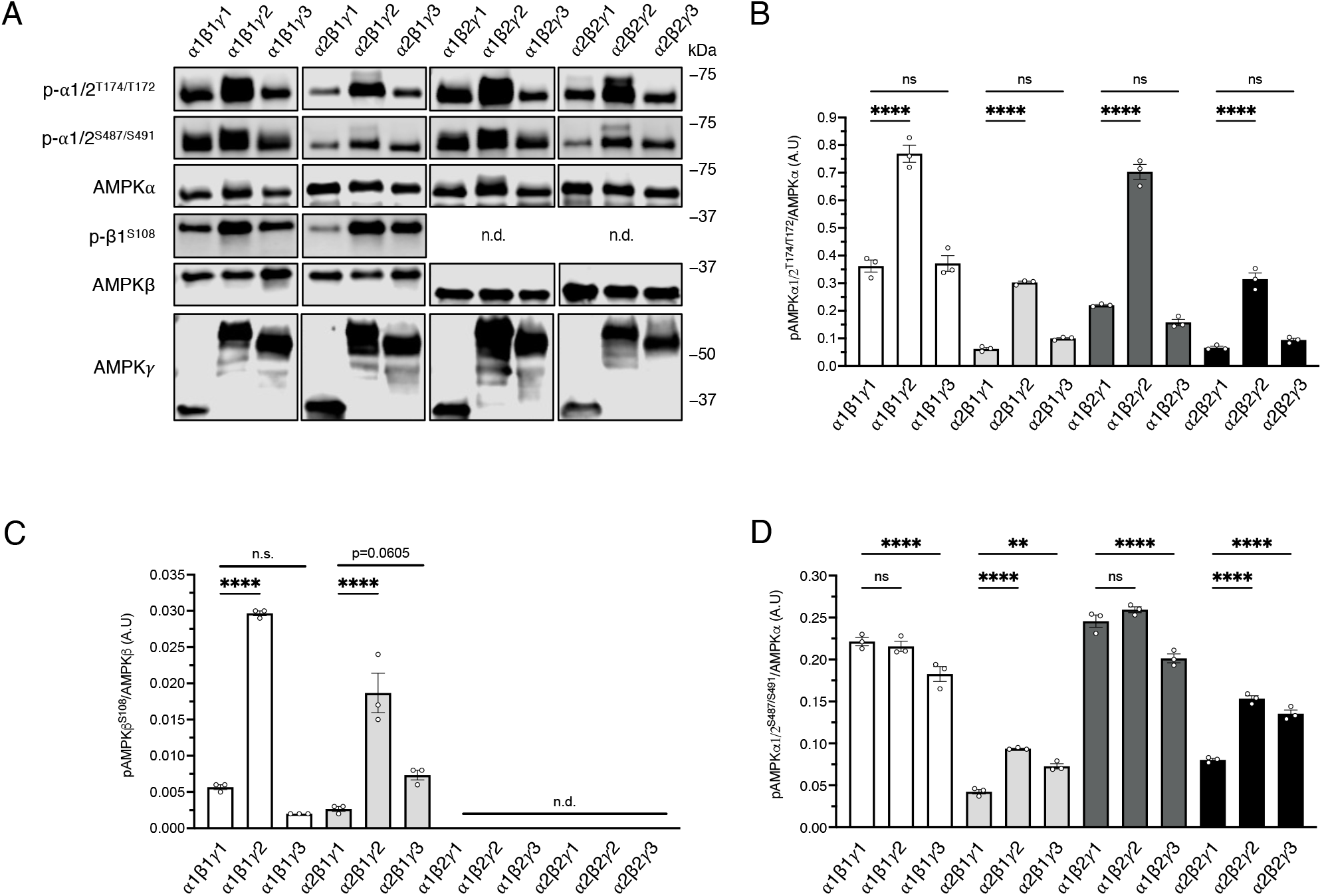
Basal comparison of regulatory phosphosites across all 12 AMPK complexes. **A** 12 AMPK complexes were FLAG-immunoprecipitated from HEK293T/17 cells incubated in complete growth media and immunoblotted as indicated. Representative immunoblots from 3 independent experiments are shown. Quantitation of **B** p-α^T172^, **C** p-β1^S108^, and **D** p-α^S487^. Data presented as average phosphorylation (arbitrary units) ± SEM, n = 3. Statistical analyses were performed by one-way ANOVA with Dunnett’s multiple-comparisons test. **P<0.01, ****P<0.0001 vs. respective γ1 complex. n.s. not significant; n.d. not determined.

**Figure 2.**
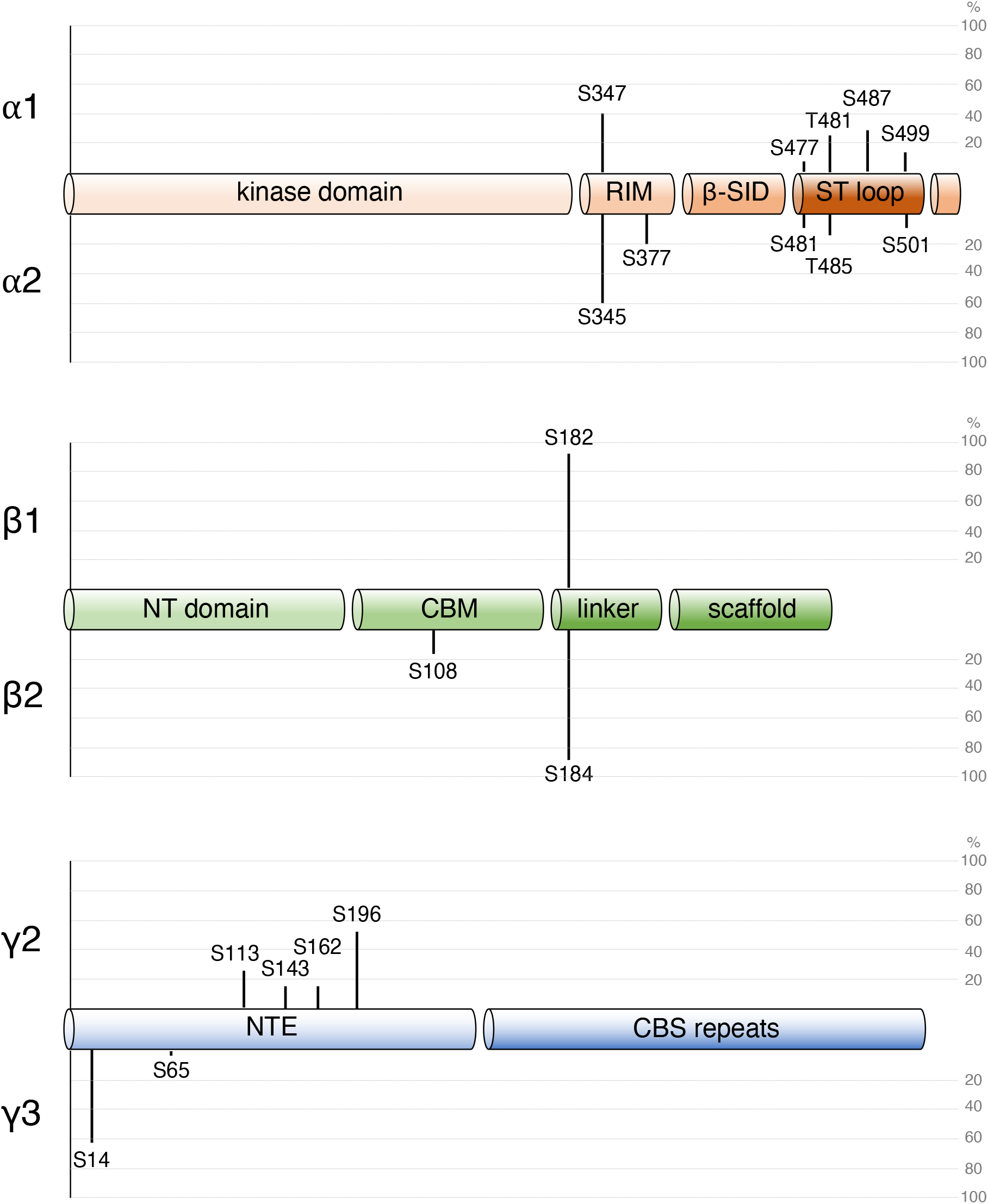
MS-detected AMPK phosphosites by subunit domain. 12 AMPK complexes were FLAG-immunoprecipitated from HEK293T/17 cells incubated in complete growth media and subjected to tryptic digest. Peptides and phosphopeptides were detected by LC-MS and area under the curve was used to calculate stoichiometries. Lengths of each marker line represent average % stoichiometry across all four complexes containing each isoform. RIM: regulatory subunit-interacting motif; -SID: -subunit interacting domain; ST loop: Ser/Thr-rich loop; NT: NH_2_-terminal; CBM: carbohydrate binding module; NTE: NH_2_-terminal extension; CBS: cystathionine β-synthase.

β1^S108^ was originally shown to be sub-stoichiometrically phosphorylated in rat liver AMPK preparations [32], and we later estimated by immunoblot that β1^S108^ was ∼6% phosphorylated in GST-tagged α1β1γ1 overexpressed in COS7 cells [20]. In the present study, p-β1^S108^ could not be accurately quantified by MS due to the presence of an interfering mass species eluting at a similar retention time. We were, however, able to quantify the corresponding p-β2^S108^ stoichiometry, albeit at low levels (4-5%) when complexed with γ1 (Table 1). Notwithstanding the co-eluting contaminant, from the MS data we could at least approximate the level of β1^S108^ phosphorylation, which is also a substrate of the autophagy initiator ULK1 [33], to be 5-10% within γ1-AMPK (data not shown). As an autophosphorylation site, it was unsurprising that β^S108^ phosphorylation was higher within γ2 complexes (Figure 1C & Table 1) that have elevated p-α^T172^ and thus higher AMPK activity.

### ST loop phosphorylation profiles

Our MS coverage of AMPK resulted in appreciable detection of several phosphorylation sites situated in the highly regulated ST loop. Firstly, we were able to quantify p-α1^S487^ (but not α2^S491^; hereafter both sites will be referred to as α^S491^) at stoichiometries in the order of 20-30% depending on γ isoform (Table 1), with lowest stoichiometries detected in γ3-AMPK. α1^S487^ is a substrate of kinases in the AGC family such as Akt [12], whereas the equivalent α2^S491^ is an autophosphorylation site [34,35]. The inability to detect p-α2^S491^ by MS appears to stem from the α2-specifc residue P498 blocking tryptic cleavage, as we can detect the analogous mouse α2 peptide harbouring a natural P498A, which is approximately 20% phosphorylated on γ1-AMPK (data not shown). Although we could not quantify human p-α2^S491^ by MS, it was assessed alongside p-α1^S487^ by immunoblot using a phospho-antibody that detects both sites (Figure 1D). The immunoblot data support our MS data suggesting γ3-AMPK complexes have slightly lower levels of p-α1^S487^. Conversely, it is clear γ2- and γ3-containing complexes have higher levels of phosphorylation on α2^S491^. While it is tempting to speculate from our immunoblot data that α1^S487^ is more highly phosphorylated than α2^S491^ we cannot rule out differences in antibody binding affinity, which is likely since mouse α2^S491^ is phosphorylated to a similar degree to human α1^S487^.

In addition, we also detected previously identified GSK3β phosphorylation sites situated on the ST loop, p-α1^S477^/α2^S481^ (p-α^S477^) and p-α1^T481^/α2^T485^ (p-α^T481^), which are intrinsically connected as p-α1^T481^ is a traditional GSK3β priming site for α^S477^ phosphorylation [36]. Basal levels of p-α^S477^ were relatively consistent across different AMPK combinations (4-18%) with stoichiometries again higher in γ2-AMPK, and in particular α2γ2-complexes (Table 1). p-α^T481^ levels (8-32%) were generally higher in α1-AMPK and highest when complexed with γ2. Interestingly, disparities between levels of p-α^S477^ and p-α^T481^ were greatest in γ3-complexes, suggesting that although effectively primed, they may be less susceptible to negative regulation by GSK3.

Within the ST loop we identified two previously uncharacterised sites, α1^S499^ and α2^S501^, where p-α1^S499^ levels (8-17%) were lowest in γ1-AMPK, and p-α2^S501^ levels (6-10%) were consistent between AMPK heterotrimers. α1^S499^ was previously shown to be phosphorylated *in vitro* by PKA [37], however it is flanked by several acidic residues that point to a casein kinase 2 recognition motif [38]. The sequence surrounding α2^S501^ (RPRSS^501^FDST), however, appears to be a better fit for a PKA consensus motif (R/K-R/K-X-S/T, where X is any amino acid) [39]. The effect of ST loop phosphorylation is generally considered to limit AMPK activity by reducing net α^T172^ phosphorylation [34–36,40–43]. Since γ2 complexes possessed both the highest level of ST loop and α^T172^ phosphorylation, this suggests the effect of attenuating p-α^T172^ may be context-dependent, or the γ2-NTE effect on p-α^T172^ protection dominates.

### Ser-Pro and γ-subunit phosphorylation site profiles

Our analysis detected several Ser-Pro phosphorylation sites. In addition to α1^S347^/α2^S345^ (α^S345^), an mTORC1 substrate that negatively regulates AMPK signalling [27], we also identified α2^S377^, a reported CDK1 substrate involved in mitotic progression [44], and β1^S182^/β2^S184^ (β^S182^), which has been shown to regulate nuclear trafficking of AMPK [45]. Additionally, the γ2-NTE contains a total of 14 Ser-Pro motifs and we managed to detect seven phosphorylation sites in this region with at least four containing that motif (due to low levels of phosphorylation of 1-3% we were unable to determine the exact phosphosite for three phospho-peptides).

α2^S345^ phosphorylation (59-70%) was considerably higher than α1^S347^ (34-45%), with both exhibiting only minor differences between β- and γ-subunit combinations (Table 1). α2^S377^ was comparably (15-22%) phosphorylated in γ1 and γ2 complexes, with slightly higher stoichiometries in γ3-AMPK (24-27%). We were able to detect the dephosphorylated peptide containing the α1-equivalent residue T373, yet basal phosphorylation was not evident despite good conservation in the consensus motif. Of all the sites identified, β^S182^ was clearly phosphorylated to the highest degree, approaching 100% stoichiometry within some of the γ1-containing complexes. β^S182^ was generally phosphorylated to a lesser degree in α2β2-complexes, although stoichiometry remained at or above 80 %. The β^S182^ data here corroborates previous work demonstrating β1^S182^ is stoichiometrically phosphorylated in largely α1-AMPK extracted from rat liver [32], whereas in rat skeletal muscle, in which α2β2 expression predominates, p-β2^S184^ is sub-stoichiometric [46]. The exclusive expression profile of γ3 in skeletal muscle suggests this tissue would give rise to AMPK heterotrimers with the highest turnover of p-β2^S184^.

Many of the γ2-NTE phosphorylation sites we detected have been identified in high throughput phosphoproteomic studies [47], yet none have undergone targeted analysis to be validated as legitimate sites. No differences in phosphorylation stoichiometries were discernible among the various α combinations. Of the γ2 sites, three are basally phosphorylated at <4%, and we are yet to identify the modified residue on the following phospho-peptides: ^133^ESSPNSNPATS^143^PGGIR^148^ was di-phosphorylated with one unknown site and ^156^TSGLSSS^162^PSTPTQVTK^171^ was tri-phosphorylated with two unknown sites. Of the four identifiable γ2-NTE sites (γ2^S113^, γ2^S143^, γ2^S162^ and γ2^S196^), all contain Ser-Pro motifs with appreciable levels (10-56%) of basal phospho-stoichiometry. Perhaps most noticeable is that the sequences flanking many of these γ2 sites not only bear resemblance with AMPK sites on distinct subunits, such as β^S182^ and the known mTORC1 substrate α^S345^, but also other *bona fide* mTORC1 substrates (Figure 3A) [48]. We also identified two phosphorylation sites residing on the γ3-NTE, S14 which is highly phosphorylated in the basal state (>60%), and S65 with low levels of phosphorylation (<4%), neither with discernible differences between α combinations (Table 1). Phosphorylation of γ3^S14^ or γ3^S16^ was previously reported on bacterially expressed α22γ3, however the authors were unable to determine which of the sites were modified [49]. Additionally, γ3^S65^ phosphorylation was detected in human muscle but is not regulated by exercise [50].

**Figure 3.**
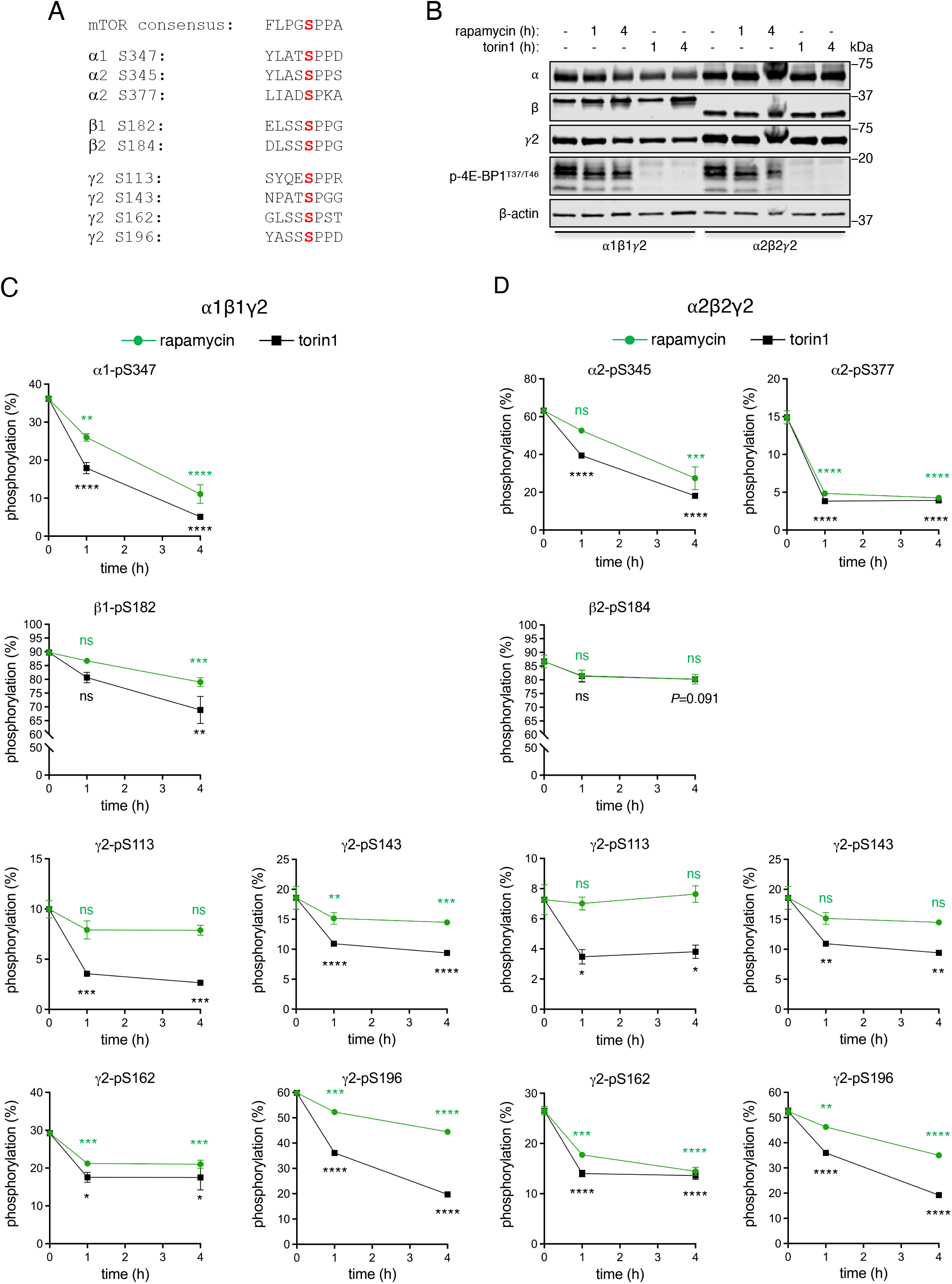
AMPK phosphosites sensitive to pharmacological inhibition of mTORC1. **A** Sequence alignment of selected Ser-Pro phosphosites on AMPK with the mTOR consensus motif [47]. **B** HEK293T/17 cells expressing α1β1γ2 or α2β2γ2 were treated with rapamycin (100 nM) or torin1 (1 μM) for up to 4 hours and lysates immunoblotted as indicated. Representative immunoblots from 3 independent experiments are shown. **C** α1β1γ2 and **D** α2β2γ2 complexes from **B** were FLAG-immunoprecipitated and subjected to tryptic digest. Peptides and phosphopeptides were detected by LC-MS and area under the curve was used to calculate stoichiometries, rapamycin and/or torin1 sensitive phosphosites at 1 h and 4 h post-treatment are depicted here. Data presented as average % stoichiometry ± SEM, n = 3. Statistical analyses were performed by one-way ANOVA with Dunnett’s multiple-comparisons test. *P<0.05, **P<0.01, ***P<0.001, ****P<0.0001 vs. basal. n.s. not significant.

### Validation of mTORC1 phosphorylation sites

Given the abundance of Ser-Pro phospho-sites across the various AMPK subunits, we speculated that some might be mTORC1 substrates. We treated HEK293T/17 cells expressing α1β1γ2 or α2β2γ2 with rapamycin, a mild mTORC1-specific inhibitor, or torin1, a more efficient ATP-competitive inhibitor of mTOR (Figure 3B) [51]. As expected, both inhibitors elicited dephosphorylation of α^S345^, but additionally, α2^S377^, β^S182^, γ2^S162^ and γ2^S196^ (Figure 3C & D). However, γ2^S113^ and γ2^S143^ phosphorylation levels were sensitive to torin1 exposure but resistant to rapamycin, a feature of several mTORC1 substrates since rapamycin only partially inhibits mTORC1 signalling (Figure 3B-C) [51].

Taking advantage of a commercially available phospho-specific antibody, β^S182^ was confirmed as an mTORC1 substrate *in vitro* using purified enzymes (Figure 4A). mTORC1 efficiently phosphorylated β^S182^, regardless of the β-isoform, in kinase dead α1-AMPK, but with far less efficiency when the β-subunit was complexed to kinase dead α2-AMPK. α isoform-specific phosphorylation of β^S182^ was confined to the β-subunit, as mTORC1 phosphorylated α^S345^ with comparable kinetics in all 4 complexes tested. Given the poor sequence conservation in the ST loops of α1 and α2, we suspected it might be responsible for the preference for α1 shown by mTORC1 when phosphorylating β^S182^. In support of this hypothesis, exchange of the α2 ST loop for the α1 sequence in α2-AMPK (α2-α1 ST loop chimera) was sufficient to recover *in vitro* mTORC1 phosphorylation of β1^S182^ on AMPK expressed in bacteria (Figure 4B). We expressed α1, α2 or the α2-α1 ST loop chimera with 2 and γ3 in HEK293T/17 cells (Figure 4C) to test if this region also influenced phosphorylation of β^S182^ *in cellulo*. Subsequent analysis demonstrated the ST loop exchange fully reintroduced α1 ST loop phosphorylation sites into the α2 construct with all all stoichiometries equivalent to α1 WT, except for the minor site p-α1^S477^ (Figure 4D & H), yet without affecting p-α^T172^ levels (Figure 4E). Notably, the α2-α1 ST loop chimera also displayed phosphorylation of β2^S184^ at levels comparable to that of α1β2γ3, as assessed by immunoblot (Figure 4F) and MS (Figure 4G). Combined, our data indicate that a region within the α1-subunit ST loop promotes accessibility of β^S182^ to mTORC1, whereas γ2 and γ3 NTEs increase the turnover of β^S182^ phosphorylation in mammalian cells.

**Figure 4.**
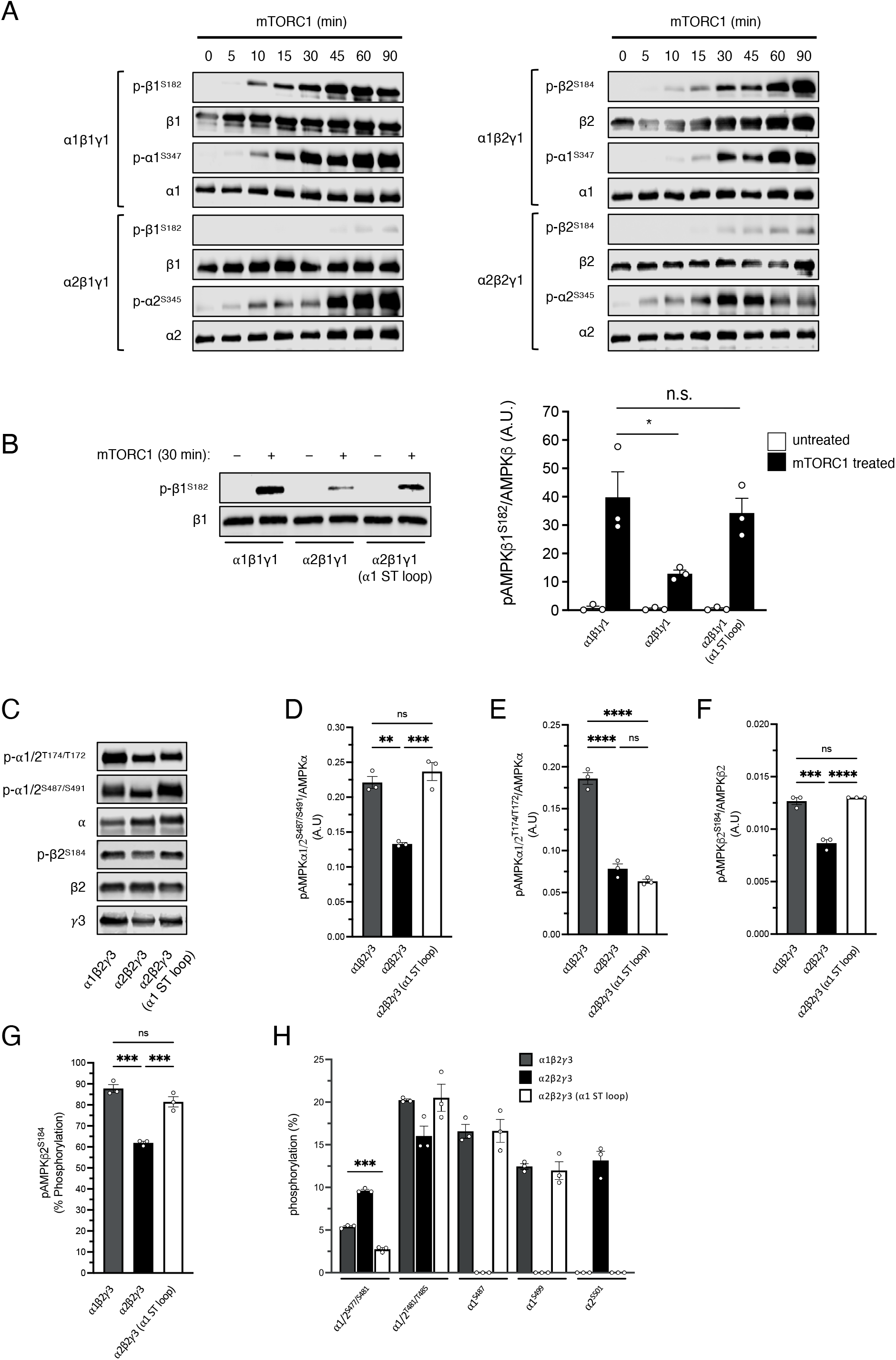
The AMPK α2-ST loop blocks mTORC1 phosphorylation of β^S182^ on purified AMPK and is associated with reduced cellular p-β2^S184^. **A** All four γ1-AMPK complexes were purified from *E. coli* as kinase dead mutants, phosphorylated by mTORC1 *in vitro* for up to 90 min and immunoblotted as indicated. **B** β1γ1, complexed to α1, α2 or α2 containing the α1 ST loop sequence (α2-α1 ST loop chimera) were purified from *E. coli* as kinase dead mutants, phosphorylated by mTORC1 *in vitro* for 30 min and p-β1^S182^ measured by immunoblot. Data presented as average β1^S182^ phosphorylation ± SEM, n = 3. Statistical analyses were performed by one-way ANOVA with Dunnett’s multiple-comparisons test. *P<0.05. n.s. not significant. **C** Lysates were prepared from HEK293T/17 cells expressing β2γ3 complexed to α1, α2 or α2-α1 ST loop chimera, AMPK complexes were FLAG-immunoprecipitated and immunoblotted as indicated. Quantitation of **D** p-α^S487^, **E** p-α^T172^, and **F** p-β2^S184^ measured by immunoblot. Data presented as average phosphorylation (arbitrary units) ± SEM, n = 3. Statistical analyses were performed by one-way ANOVA with Dunnett’s multiple-comparisons test. **P<0.01, ***P<0.001, ****P<0.0001. n.s. not significant. AMPK complexes from **C** were subjected to tryptic digest. Peptides and phosphopeptides were detected by LC-MS and area under the curve used to calculate stoichiometries of **G p-**β2^S184^ and **H** ST loop phosphosites. Data presented as average % stoichiometry ± SEM, n = 3. Statistical analyses were performed by one-way ANOVA with Dunnett’s multiple-comparisons test. ***P<0.001. n.s. not significant. Representative immunoblots from 3 independent experiments are shown.

### Regulation of β^S182^ in mammalian cells

To gain a better understanding of p-β^S182^ regulation in mammalian cells, HEK293T cells expressing γ1 with each α isoform combination (1/2 and FLAG-α1/2), were treated with 250 nM torin1 for an extended time-course up to 24 h, and lysates or FLAG-immunoprecipitated AMPK analysed by immunoblotting and MS. Consistent with the shorter torin1 incubations (Figure 3B-D), dephosphorylation of β1^S182^ and β2^S184^ occurred gradually over time in both α1- and α2-AMPK up to 8 h (Figure 5A-D). By immunoblot we detected a smaller degree of β2^S184^ dephosphorylation post-8h for both α-isoforms, however in all complexes even at 24 h, ∼50% of basal signals were retained. Similar trends were detected using MS to measure absolute stoichiometry, with dephosphorylation of β1^S182^ andβ2^S184^ bottoming out at 8h after torin1 addition, and 65-75% of respective total β1 or β2 subunits remaining phosphorylated at these sites. 24 h incubation with rapamycin (1 μM) did not substantially affect p-β^S182^ or p-β2^S184^ levels, and torin1 incubation (1 μM) was unable to induce complete dephosphorylation of p-β1^S182^, on endogenous AMPK in HEK293T cells (Figure 5E), indicating the presence of torin1 insensitive signalling pathways contributing to β1^S182^ phosphorylation. Unlike the effect of pharmacological mTORC1 inhibition, classical activators of AMPK that disrupt mitochondrial ATP production (phenformin, H_2_O_2_) or glycolysis (2-DG) had no bearing on levels of phosphorylation of β^S182^ on AMPK expressed in COS7 cells (Figure 5F & G).

**Figure 5.**
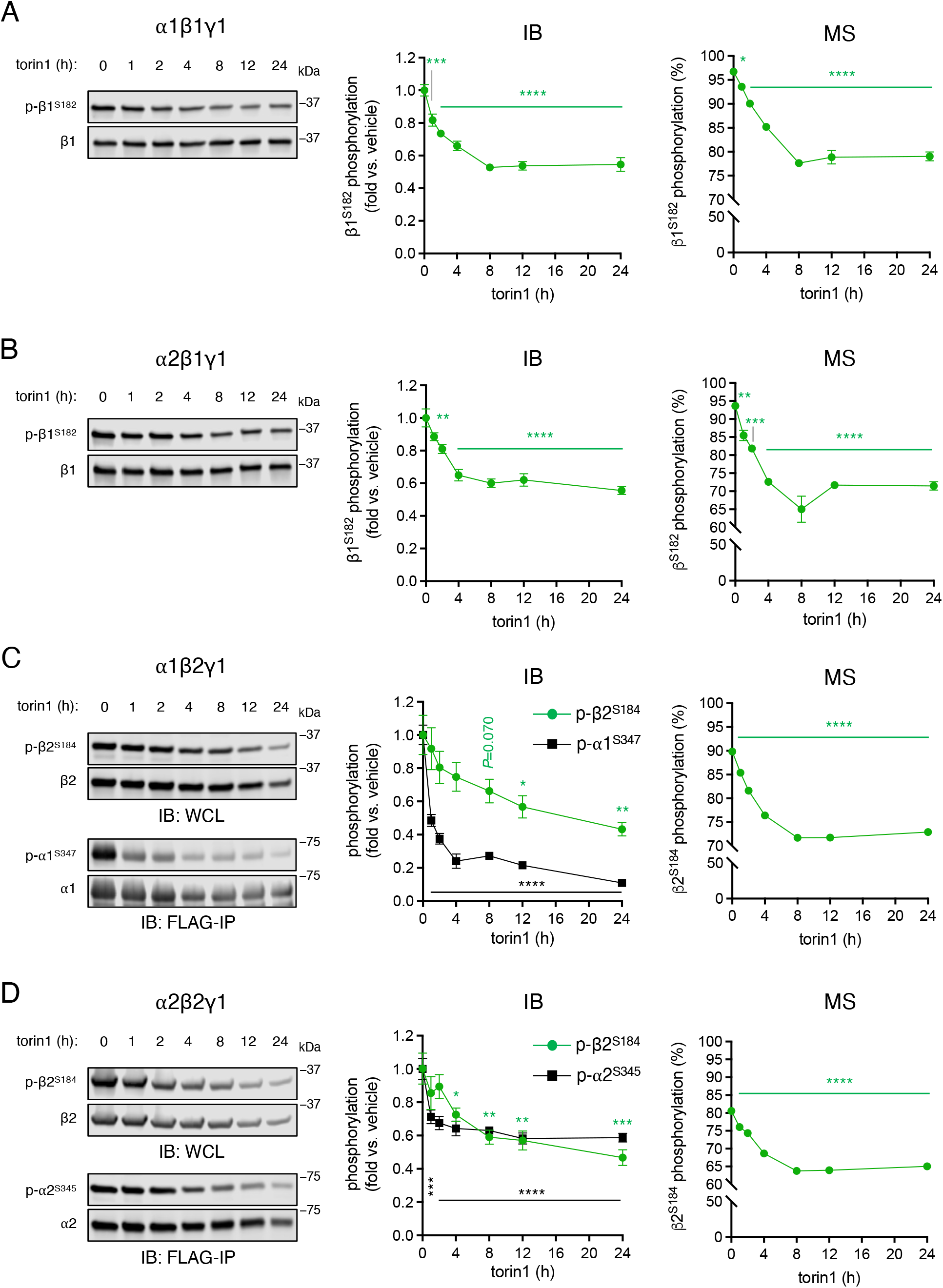
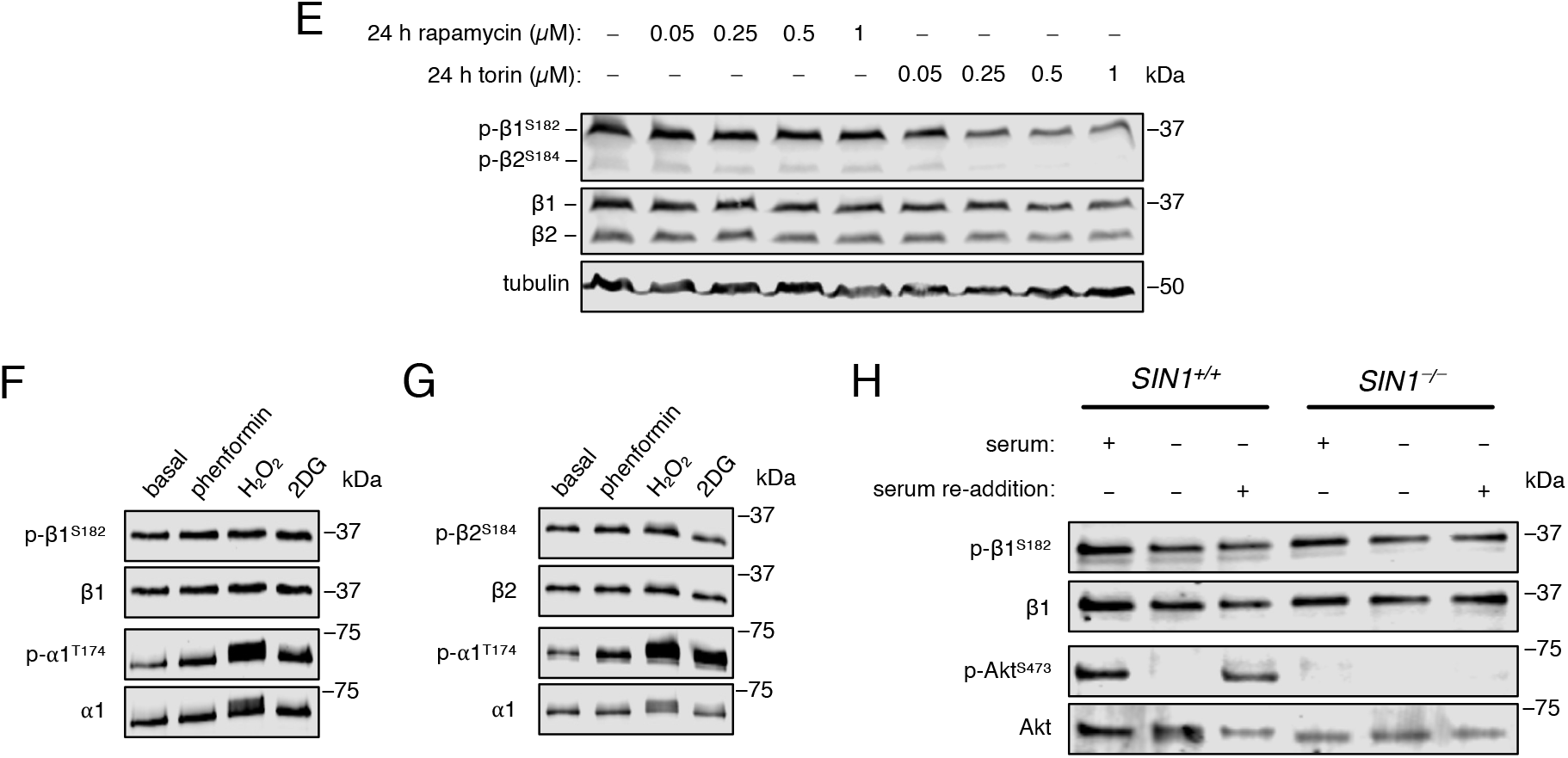
Extended mTORC1 inhibition by torin1 leads to partial dephosphorylation of p-β^S182^ in HEK293T cells. HEK293T cells expressing FLAG-fusions of **A** α1β1γ1, **B** α2β1γ1, **C** α1β2γ1, or **D** α2β2γ1 were incubated with 250 nM torin1 for up to 24 h, and indicated phosphosites/stoichiometries detected from prepared lysates or FLAG-immunoprecipitated AMPK by immunoblot or MS. Data presented as average phosphorylation (arbitrary units) ± SEM, or average % stoichiometry ± SEM, n = 3. Statistical analyses were performed by one-way ANOVA with Dunnett’s multiple-comparisons test. *P<0.05, **P<0.01, ***P<0.001 and ****P<0.0001 vs. basal. **E** Lysates were prepared from HEK293T cells incubated with rapamycin or torin1 (0.05 to 1 μM) for 24 h and immunoblotted for endogenous p-β^S182^. COS7 cells expressing GST-fusions of **F** α11γ1 or **G** α1β2γ1 were incubated with phenformin (2 mM, 1 h), H_2_O_2_ (5 mM, 45 min) or 2-deoxy-glucose (2DG; 25 mM, 0.5 h), and GST-purified AMPK was immunoblotted as indicated. **H** Lysates were prepared from immortalised MEFs (WT or SIN1^-/-^), incubated under conditions of serum starvation overnight ± subsequent serum re-addition (1 h), and immunoblotted as indicated. Representative immunoblots from 3 independent experiments are shown.

mTORC2 is known to phosphorylate turn motif sites in AGC kinases like Akt during translation to ensure stability of the polypeptide upon release from the ribosome [52,53]. To investigate whether β^S182^ is an mTORC2 substrate in mammalian cells, iMEFs genetically devoid of the critical mTORC2 component SIN1 were either cultured in complete media, or nutrient starved and subsequently restimulated by adding back nutrient-replete media. Treatment efficacy was confirmed by loss (nutrient starved) and recovery (nutrient replenishment) of mTORC2-catalysed Akt^S473^ phosphorylation, which was completely absent in SIN1-KO cells (Figure 5H). However, there were no differences in p-β1^S182^ between genotypes or in response to the treatments.

### Revisiting functional roles for β^S182^ phosphorylation

β^S182^ was first identified as a phosphorylation site in 1997 [32]. Subsequent analyses using the β1-S182A mutant revealed that phosphorylation at this site did not affect basal activity of AMPK complexes, nor sensitivity to AMP [45], which here we confirm for both β1^S182^ and β2^S184^ (Figure 6A-F). Given the proposed structural proximity of β^S182^ to the AMPK ADaM site, we speculated phosphorylation at this site may influence drug sensitivity. However, EC_50_ values for the ADaM drug A-769662 were comparable between WT and β1-S182A mutant AMPK isolated from COS7 cells when assayed *in vitro*, and maximal fold activation was unchanged (Figure 6C). Additionally, the half-lives of WT β1 and the β1-S182A mutant were virtually identical in HEK293Ts treated with cycloheximide for 12 h (Figure 6G), an inhibitor of protein synthesis, demonstrating this site is not required to stabilise the AMPK heterotrimer.

**Figure 6.**
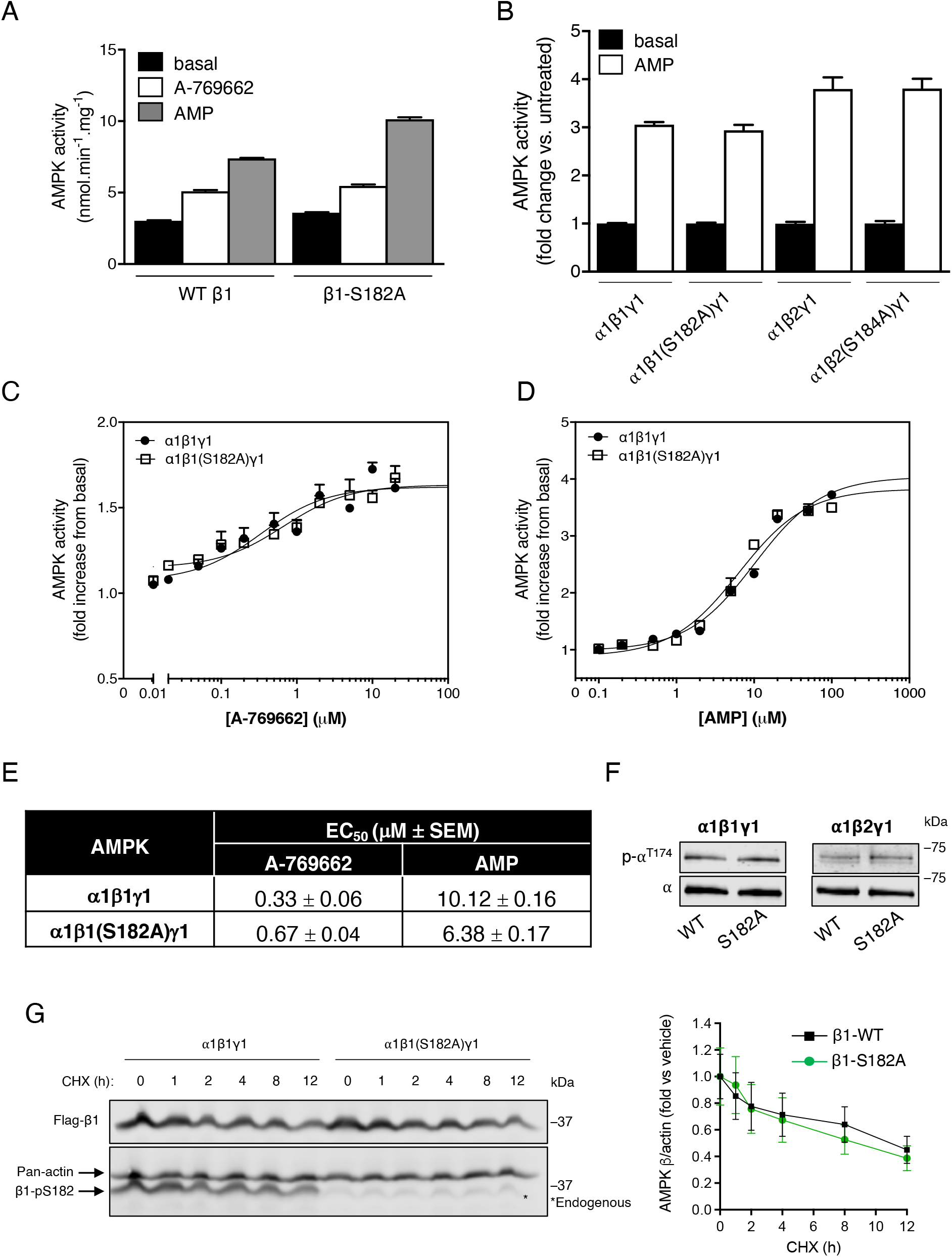
Investigating regulatory functions of p-β^S182^. **A** Sensitivities of AMPK α1β1γ1 (WT or 1-S182A mutant), FLAG-purified from COS7 cells incubated in complete growth media, to AMP (100 μM) or A-769662 (20 μM) were determined by radiolabelling of SAMS peptide substrate. Data presented as average AMPK activity (nmol.min^-1^.mg^-1^) ± SEM, n = 3. Statistical significance was calculated by one-way ANOVA with Dunnett’s multiple-comparisons test. *P<0.05, **P<0.01, ***P<0.001, ****P<0.0001. **B** Sensitivities of AMPK α1β1γ1 or α1β2γ1 (WT or respective -S182A mutants), FLAG-purified from COS7 cells incubated in complete growth media, to AMP (100 μM) were determined by radiolabelling of SAMS peptide substrate. Data presented as average fold change in AMPK activity (nmol.min^-1^.mg^-1^) vs. basal ± SEM, n = 3. Statistical analyses were performed by unpaired t test. *P<0.05, **P<0.01, ***P<0.001, ****P<0.0001. Activities of FLAG-purified AMPK α1β1γ1 (WT or β1-S182A mutant) to increasing doses of **C** A-769662 or **D** AMP were determined by radiolabelling of SAMS peptide substrate. Data presented as average fold change in AMPK activity (nmol.min^-1^.mg^-1^) vs. basal ± SEM, n = 4. **E** EC_50_ values for A-769662 and AMP activation of α1β1γ1 and β1-S182A mutant, calculated from dose curves in (C) and (D). **F** Lysates were prepared from COS7 cells, expressing either AMPK α1β1γ1 or α1β2γ1 (WT or respective β-S182A mutants) and incubated in complete growth media, and immunoblotted for p-α^T172^. **G** Lysates were prepared from cycloheximide-treated (CHX; 1 μM for up to 12 h) HEK293T cells, expressing AMPK α1β1γ1 (WT or 1-S182A), and immunoblotted for total AMPK 1 subunit. Data presented as average fold change in β1/actin protein content vs. untreated ± SEM, n = 3. Statistical analyses were performed by unpaired t test. Representative immunoblots from 3 independent experiments are shown.

β^S182^ phosphorylation has been proposed to negatively regulate nuclear translocation of AMPK in HEK293 cells [45]. Despite this, we found that p-β2^S184^ dephosphorylation was not a prerequisite for AMPK nuclear localisation in HEK293T cells, as the basal phosphorylation stoichiometry of β2^S184^ complexed with α1 and γ1 was near-maximal in both nuclear and cytoplasmic compartments (Figure 7A). Torin1-induced p-β^S182^ dephosphorylation had little-to-no effect on enhancing nuclear α2β1γ1 or α2β2γ1 abundance, even at 24 h of drug exposure when residual phospho-signals were lowest (Figure 7B-D). Nevertheless, consistent with previous findings [54,55], HEK293T cells expressing 2-AMPK had increased proportional abundance of nuclear AMPK versus those expressing β1, even though both β1 and β2 were co-expressed with α2 which contains a nuclear localisation signal [54]. Collectively, our findings indicate that mTORC1 is one of potentially several upstream kinases for β^S182^, that p-β^S182^-mediated regulation of nuclear translocation may be contextual, and that phosphorylation of this site does not appear to influence allosteric (i.e., small molecule-induced) activation of AMPK.

**Figure 7.**
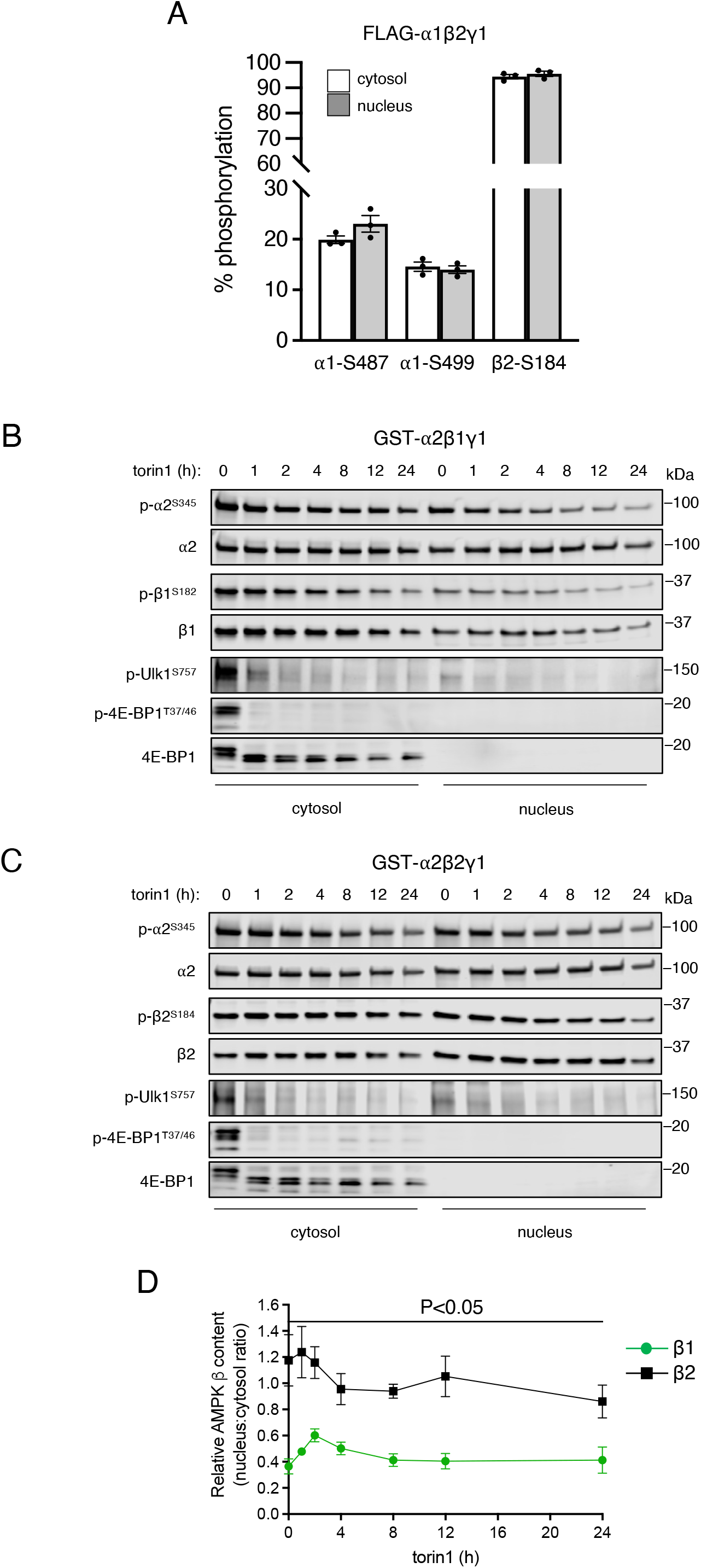
Dephosphorylation of p-β^S182^ is not a prerequisite for AMPK nuclear transport. Cytosolic and nuclear fractions were prepared from HEK293T cells expressing α1β2γ1, α2β1γ1 or α2β2γ1 and incubated in complete media. **A** Stoichiometries of p-α1^S487^, p-α1^S499^ and p-β2^S184^ on FLAG-α1β2γ1 in cytosolic and nuclear fractions determined by MS. Data presented as average % stoichiometry ± SEM, n = 3. Statistical analyses were performed by unpaired t test. Cytosolic and nuclear fractions from **B** GST-α2β1γ1 and **C** GST-α2β2γ1 expressing HEK293T cells, incubated with torin1 (250 nM) for up to 24 h, were immunoblotted as indicated, and **D** quantitated. Data are presented as relative AMPK β content (nucleus:cytosol ratio) ± SEM, n = 4. Statistical analyses were performed by multiple unpaired t test vs β1 expressing cells. Representative immunoblots from 3 independent experiments are shown.

### Dephosphorylation of β2^S184^, but not β1^S182^, enhances cell proliferation

Analysis of cell proliferation in real time by live-cell imaging in the Incucyte® system revealed the β2-S184A mutant (expressed alongside α2 and γ1) enhanced the growth of HEK293 cells relative to WT 2 in response to amino acid (Arg, Lys) withdrawal (Figure 8A & B). This effect was specific to β2-containing complexes, as no such differences in growth were observed for 1-expressing cells. We therefore posited that β2^S184^ phosphorylation could occupy a more prominent role in regulating nuclear AMPK activity, rather than trafficking. To address this, *β1/2-*double knockout iMEFs were introduced, via lentiviral-mediated delivery, with FLAG-tagged human AMPK β2 subunits (WT or β2-S184A) to reconstitute the AMPK heterotrimer [33]. Indeed, nuclear AMPK in β2-expressing iMEFs had higher specific activity than its cytoplasmic counterpart, an effect enhanced by the β2-S184A mutant (Figure 8C). It is interesting that increased 2-S184A-induced nuclear AMPK activity correlated with enhanced cell proliferation in response to nutrient stress, in contrast to delayed proliferation of cells expressing the activating α2-S345A mutant under identical experimental conditions [27].

**Figure 8.**
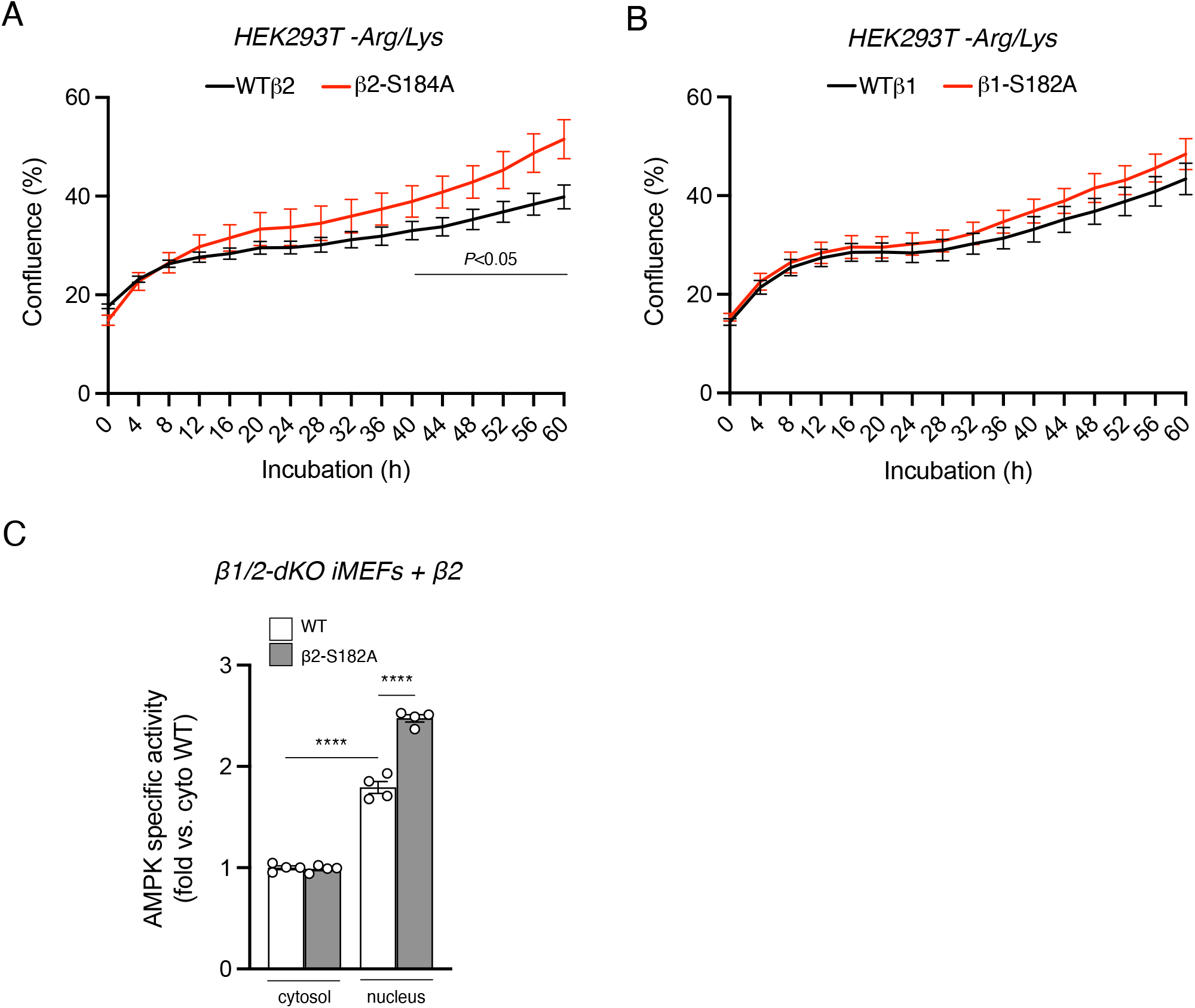
Dephosphorylation of p-β2^S184^ promotes AMPK nuclear activity and cell proliferation under conditions of amino acid stress. Real-time proliferation analysis of HEK293T cells expressing GFP-fusions of either **A** α1β2γ1 (WT or 2-S184A mutant) or **B** α1β1γ1 (WT or β1-S182A mutant), after switching to arginine- and lysine-free medium from complete growth medium. Data presented as average % confluence ± SEM, n = 3. Statistical analyses at each time point were performed by multiple unpaired t test vs. respective WT. **C** Cytosolic and nuclear fractions were prepared from β1/2-dKO iMEFs expressing 2 WT or S184A mutant, and activities of FLAG-immunoprecipitated AMPK measured by radiolabelling of SAMS synthetic peptide. Data presented as fold change in average AMPK specific activity vs. cytosolic WT ± SEM, n = 3. Statistical analyses were performed by unpaired t test. ****P<0.0001.

## Discussion

In the present study we have taken advantage of a targeted MS approach that, coupled with conventional immunoblotting, generated comprehensive phosphorylation profiles of all 12 human AMPK complexes expressed in mammalian cells. AMPK is subject to a multitude of posttranslational modifications, most notably phosphorylation [12], with canonical activation occurring via p-α^T172^ by LKB1 and CaMKK2. Here, under optimal cell growth conditions we demonstrate substantial heterogeneity in AMPK phosphorylation of α^T172^, β^S108^ (which sensitises AMPK to ADaM site ligands) and a range of α-subunit ST loop sites (several of which thought to limit p-α^T172^ [34–36,40–43]). Moreover, of the 18 phosphorylation sites our analysis detected, 13 were of previously unknown function, with several displaying sensitivity to pharmacological inhibition of mTOR. This included β^S182^, which is by far the most heavily phosphorylated of all AMPK sites that at times approximated 100% stoichiometry, in line with other studies [45].

We discovered β^S182^ is a direct substrate of mTORC1, in which the presence of α1 in the heterotrimer augmented mTORC1 phosphorylation of β^S182^ in a ST loop-dependent manner, a region that diverges between the two α isoforms. Whether phosphorylation of an α1 ST loop phosphosite(s) promotes, or phosphorylation of an α2 ST loop phosphosite(s) inhibits, the mTORC1 β-subunit interaction is an area for future interrogation. Nevertheless, it is worth highlighting that Akt, which in response to growth factors indirectly activates mTORC1 by relieving inhibition by TSC2 [56], is an α1 ST loop kinase and potential candidate for facilitating β^S182^ phosphorylation. In agreement with previous analysis of the β2-subunit extracted from skeletal muscle [32,46], α2β2γ2 and α2β2γ3 complexes (highly expressed in this tissue) had the lowest AMPK β^S182^ phosphorylation stoichiometry, suggestive of greater rates of turnover and a functional manifestation. Indeed, dephosphorylation of β2^S184^ preferentially upregulated nuclear AMPK activity and enhanced cell proliferation in the face of nutrient restriction. In skeletal muscle cells, simply exchanging β2 with β1 in complex with α2 (that contains a nuclear localisation signal) renders AMPK refractory to nuclear translocation [54], while in rat liver α1-AMPK infiltrates the nucleus in a circadian-dependent manner commensurate with peak expression of β2 and not β1 [55]. The myristoylation-deficient β1-G2A mutant, that abolishes membrane-bound organelle (i.e., lysosome) association, can, however, translocate to the nucleus upon stimulation [54]. Whilst we have not definitively confirmed whether p-β^S182^ dephosphorylation has a bearing on AMPK nuclear localisation [45], it does appear that the intrinsic properties of the β2 isoform are critical determinants of nuclear AMPK biology.

Our findings may reconcile, in part, the complex role AMPK plays in cancer [57], whereby in the established tumour, AMPK endows cancerous cells with an ability to proliferate despite the poorly vascularised and nutrient-scant microenvironment. TFEB is a master transcriptional regulator of autophagosome and lysosome biogenesis that ordinarily, is inhibited by mTORC1 by multisite phosphorylation [58–60]. However, TFEB is constitutively active alongside mTORC1 in some cancers such as Birt–Hogg–Dubé syndrome, a disease caused by loss-of-function mutations in *FLCN* that leads to renal cell carcinoma [61]. TFEB was recently shown to be a substrate of AMPK [62], in which inhibition of its phosphorylation by AMPK attenuated chemoresistance. Whether mTORC1 inhibition-induced p-β2^S184^ dephosphorylation, in turn prioritising nuclear AMPK activity and substrate (e.g., TFEB) targeting, contributes to chemoresistance is an area of future research. In addition to the well-characterised RAG-GTPase-dependent scaffolding of mTORC1 to the lysosome for activation [21–24], there are newly discovered, distinct modes of mTORC1 activation in the nucleus [63,64]. As previously mentioned, β2^S184^ dephosphorylation drives cell proliferation during nutrient stress (presumably due to upregulated autophagy [27]); however, dephosphorylation of another mTORC1 substrate on AMPK, α2^S345^, has the opposite effect under identical conditions [27]. Thus, there may exist compartment-specific modes of mTORC1 and AMPK signalling to fine-tune prevailing cellular responses to changes in nutrient availability. This is potentially made possible by the existence of up to 12 unique AMPK heterotrimers that are each differentially regulated by phosphorylation.

In conclusion, our unique phosphorylation profiles of AMPK will serve as a useful roadmap for future studies focussing on its regulation in cells and tissues, where preferential isoform expression patterns influence entire signalling networks, such as γ2, an important aspect of cardiac physiology [65–67]. Our analysis of torin1 sensitive phosphosites reveal that AMPK is extensively regulated by mTOR signalling across many of its subunits, which highlights the need for future investigations to thoroughly interrogate this signalling axis. The discoveryβ^S182^ as an mTORC1 substrate that regulates nuclear AMPK activity and cell proliferation during nutrient restriction, will help inform future work aimed at deciphering the complex actions of AMPK in cancer and disease recurrence.

## Author Contribution

BEK, JP and JSO conceived the project; WJS, AJO, DY, NXYL, KRM, AH and CGL designed and performed experiments and analysed data. WJS, AJO and DY performed LC-MS experiments; JWS & SG provided conceptual input and reagents; WJS, AJO and JSO wrote the manuscript. All authors discussed the results and approved the final version of the manuscript.

## Acknowledgements

This work was supported by grants from the National Health and Medical Research Council (no. 1161262 to JP & JSO), the Australian Research Council (no. DP170101196 to B.E.K and DP180101682 to JP.) and the Jack Brockhoff Foundation (no. JBF-4206 to C.G.L.). J.P. was supported by a Flinders Foundation seeding grant and Flinders University (Australia). C.G.L. is an NHMRC Early Career Research Fellow. This study was supported in part by the Victorian Government’s Operational Infrastructure Support Program.

## Experimental Procedures

### Materials

Rapamycin (R8781), AMP (A1752), ATP (A2383), phenformin (P7045), 2-deoxy-D-glucose (2DG; D6134), H_2_O_2_ (18304) and cycloheximide (C7698) were from Sigma-Aldrich. Torin1 (S2827) was from Selleckchem and A769662 (ab120335) was from Abcam.

### Plasmid constructs and mutagenesis

For heterotrimeric bacterial expression of the α2-α1 ST loop chimera (α2_1-474_ /α1_471-530_/α2_533-552_), the human DNA sequence for was generated and cloned into pET21b using BamHI/NotI restriction sites by Gene Universal (Newark, Delaware, United States). DNA was cloned into the pET DUET-1 multiple cloning site (MCS) 1 (MCS2 already containing γ1) using MfeI/XhoI restriction sites in-frame with an NH_2_-terminal 6xHis tag, replacing the wild-type α2 DNA sequence [20]. For mammalian expression, the α2-α1 ST loop chimera DNA was cloned into pcDNA3.1(-) using XhoI/HindIII restriction sites. For mammalian expression of γ1 and γ2, the human DNA sequences were cloned into pcDNA3.1(-) using NotI/EcoRI restriction sites. For mammalian expression of γ3, the human DNA sequence was generated and cloned into pcDNA3.1(-) using NotI/EcoRI restriction sites by Gene Universal. All primers used for cloning were designed and ordered from Sigma-Aldrich and are outlined in Table 2. All other plasmids used in this study have been described previously [17,20,33,68,69].

**Table 2.**
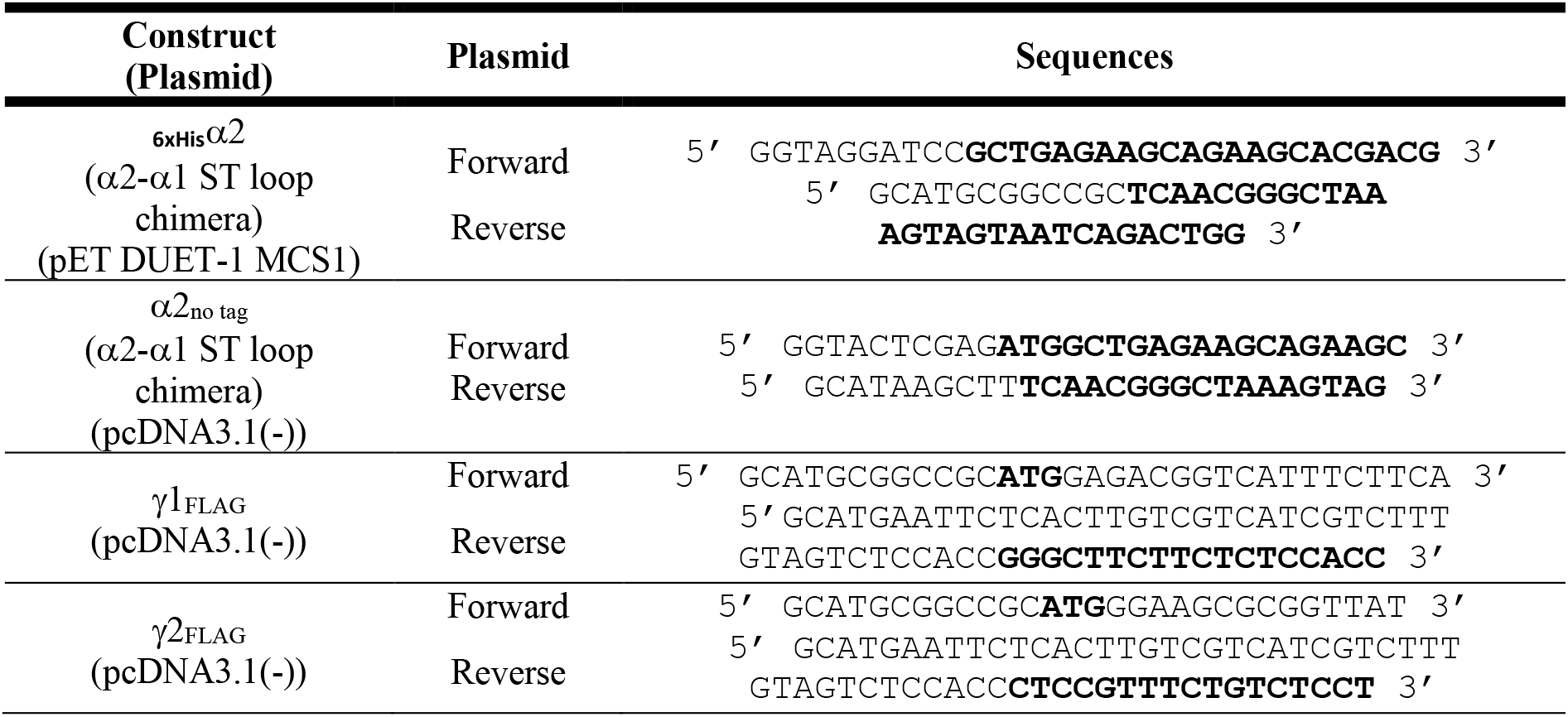
Primers used for cloning. Complimentary sequences are in bold.

For mutagenesis, primers containing the desired mutation, as detailed in Table 3, were designed and ordered from Sigma-Aldrich. Both forward and reverse mutagenic primers were mixed with 10x PfuUltra buffer (NEB), DNA template (40 ng), dNTPs (Sigma-Aldrich), PfuUltra DNA polymerase (NEB) and nuclease free H_2_O to a final volume of 50 μL. The DNA was amplified by PCR using a thermocycler (Bio-Rad). Dpn1 (NEB) was added to the reaction and incubated for 2 hours at 37 °C to digest methylated parental DNA. The amplified DNA was transformed into α-select competent cells (Bioline) by heat shock and plated on Luria-Bertani broth (LB) agar plates containing 100 μg mL^-1^ antibiotics. Individual colonies were inoculated in 5 mL LB containing 100 μgmL^-1^ antibiotics and incubated overnight at 37 °C. Plasmid DNA was isolated using a Wizard Plus SV minipreps DNA purification kit (Promega) as per the manufacturers protocol. All new constructs were confirmed by Sanger sequencing (Micromon, Monash University, Australia).

**Table 3.** Primers used for site-directed mutagenesis. Codon changes are in bold.

| Mutation | Type | Sequences |
| --- | --- | --- |
| $\beta 1$-S182A | Forward | 5' TGAGCTGTCCAGT**GCT**CCCCCAGGACCCTA 3' |
|  | Reverse | 5' TAGGGTCCTGGGGG**AGC**ACTGGACAGCTCA 3' |
| AMPK$\beta 2$-S184A | Forward | 5' CCTTTCCAGC**GCA**CCCCCAGGGC 3' |
|  | Reverse | 5' GCCCTGGGGG**TGC**GCTGGAAAGG 3' |

### Immunoblotting

Protein samples were separated on gradient SDS-PAGE gels (Bio-Rad) and transferred to an Immobilon-FL PVDF membrane (Merck Millipore). After blocking the membrane with 2 % non-fat dry milk dissolved in PBS + 0.1 % (v/v) tween-20 (PBST; Sigma-Aldrich), membranes were incubated with primary antibodies for either 2 hours at room temperature or overnight at 4 °C, as indicated in Table 4. Following repeated washes with PBST, fluorescently labelled secondary antibodies diluted in PBST were added as detailed in Table 3. The membranes were then washed extensively in PBST and visualised on the Odyssey Infrared Imaging System (LI-COR Biosciences) and immunoreactive bands analysed and quantified using Image Studio Software (LI-COR Biosciences).

**Table 4.** Antibodies used for immunoblotting

| Antibody<br>(Catalogue Number) | Type | Source | Incubation<br>Period | Dilution | Supplier |
| --- | --- | --- | --- | --- | --- |
| AMPKα<br>(2793) | Primary | Mouse | 2 hours/<br>Overnight | 1:1000 | Cell Signaling<br>Technology |
| p-AMPKα1/2<sup>T174/T172</sup><br>(2535) | Primary | Rabbit | 2 hours/<br>Overnight | 1:1000 | Cell Signaling<br>Technology |
| p-AMPKα1<sup>S347</sup><br>(In-house) | Primary | Rabbit | Overnight | 1:1000 | [27] |
| p-AMPKα2<sup>S345</sup><br>(ab129081) | Primary | Rabbit | Overnight | 1:1000 | Abcam |
| p-AMPKα1/2<sup>S487/S491</sup><br>(2185) | Primary | Rabbit | Overnight | 1:1000 | Cell Signaling<br>Technology |
| AMPKβ<br>(4150) | Primary | Rabbit | Overnight | 1:1000 | Cell Signaling<br>Technology |
| p-AMPKβ1<sup>S108</sup><br>(4181) | Primary | Rabbit | Overnight | 1:1000 | Cell Signaling<br>Technology |
| p-AMPKβ1/2<sup>S182/S184</sup><br>(4186) | Primary | Rabbit | Overnight | 1:1000 | Cell Signaling<br>Technology |
| AMPKβ<br>(4150) | Primary | Rabbit | Overnight | 1:1000 | Cell Signaling<br>Technology |
| FLAG tag<br>(9146) | Primary | Mouse | 2 hours/<br>Overnight | 1:1000 | Cell Signaling<br>Technology |
| FLAG tag<br>(14793) | Primary | Rabbit | 2 hours/<br>Overnight | 1:1000 | Cell Signaling<br>Technology |
| MYC tag<br>(2276) | Primary | Mouse | 2 hours/<br>Overnight | 1:1000 | Cell Signaling<br>Technology |
| ULK1<br>(4773) | Primary | Rabbit | Overnight | 1:1000 | Cell Signaling<br>Technology |
| p-ULK1<sup>S757</sup><br>(6888) | Primary | Rabbit | Overnight | 1:1000 | Cell Signaling<br>Technology |
| p-4E-BP1<sup>T37/T46</sup><br>(9459) | Primary | Rabbit | Overnight | 1:1000 | Cell Signaling<br>Technology |
| 4E-BP1<br>(9452) | Primary | Rabbit | Overnight | 1:1000 | Cell Signaling<br>Technology |
| p-Akt<sup>S473</sup><br>(9271) | Primary | Rabbit | Overnight | 1:1000 | Cell Signaling<br>Technology |
| Akt<br>(2920) | Primary | Mouse | Overnight | 1:1000 | Cell Signaling<br>Technology |
| β-actin<br>(4970) | Primary | Rabbit | 2 hours | 1:1000 | Cell Signaling<br>Technology |
| α-tubulin<br>(3873) | Primary | Mouse | 2 hours | 1:1000 | Cell Signaling<br>Technology |
| IRDye 680RD<br>(926-68070) | Secondary | Mouse | 1 hour | 1:10,000 | LI-COR<br>Biosciences |
| IRDye 680RD<br>(926-68071) | Secondary | Rabbit | 1 hour | 1:10,000 | LI-COR<br>Biosciences |
| IRDye 800CW<br>(926-32210) | Secondary | Mouse | 1 hour | 1:10,000 | LI-COR<br>Biosciences |
| IRDye 800CW<br>(926-32211) | Secondary | Rabbit | 1 hour | 1:10,000 | LI-COR<br>Biosciences |

### Cell culture

Generation of β1/β2 double knockout (β-dKO) iMEFs has been described previously [33]. SIN1 knockout (SIN1-KO) iMEFs were a gift from Bing Su (Shanghai Jiao Tong University School of Medicine). iMEF, HEK293T/17, HEK293T and COS7 cells (ATCC) were all maintained in Dulbecco’s Modified Eagle’s medium (DMEM; Sigma-Aldrich, D5796) supplemented with 10% fetal bovine serum (FBS; Assay Matrix) at 37 °C and 5 % CO_2_. The cultures were incubated with fresh DMEM containing 10% FBS for 2 hours prior to drug treatments.

### Mammalian protein expression and purification

For transient AMPK expression in HEK293T/17, HEK293T and COS7 cells, adherent cultures at ∼50 % confluency were triply transfected with full-length α1 or α2 (COOH-terminal FLAG fusion or untagged, pcDNA3.1 vector, or NH_2_-terminal GST fusion, pDEST27 vector), β1 or β2 (COOH-terminal MYC or FLAG fusion, pcDNA3.1 vector), and γ1, γ2 or γ3 (NH_2_-terminal HA fusion, pMT2 vector or COOH-terminal FLAG fusion, pcDNA3.1 vector) in the presence of FuGENE HD transfection reagent (Promega) for 48 hours as per the manufacturers protocol. For expression of heterotrimeric AMPK in β-dKO iMEFs, β1 or β2 (COOH-terminal FLAG fusion, LeGO-iG2 vector; WT and indicated mutants) was reintroduced by lentiviral transduction using the LeGO-iG2 system. After 24 hours, the lentivirus-containing media was replaced with fresh media, and cells were harvested 72 hours after transduction. For cell proliferation assays, HEK293 cells were seeded at ∼10-15 % confluency 48 hours before a triple transfection with Lipofectamine 2000 (Thermo Fisher Scientific) as per the manufacturers protocol.

Cells were first gently washed in ice cold phosphate buffered saline (PBS; Sigma-Aldrich) then scraped in ice-cold lysis buffer. For collection of whole cell material, cells were lysed in buffer containing 50 mM Tris pH 7.5, 150 mM NaCl, 10 % (v/v) glycerol, 50 mM NaF, 5 mM sodium pyrophosphate and 1 % (v/v) Triton X-100. Cell lysates were clarified by centrifugation (16,000 g, 3 min, 4 °C), flash frozen in L-N_2_ and stored at -80 °C until subsequent analysis. For separation of cytosolic and nuclear compartments, cells were scraped in cytosolic lysis buffer (10 mM HEPES pH 7.9, 10 mM KCl, 10 % (v/v) glycerol, 50 mM NaF, 5 mM sodium pyrophosphate, 0.1 mM EDTA, 1.5 mM MgCl_2_, 0.5 mM 1,4-dithiothreitol (DTT), 0.5 % (v/v) NP40). Lysates were incubated on ice for 10 min, centrifuged (5,000 g, 10 min, 4 °C) with the supernatant removed as the fraction containing cytosolic proteins. The pellet was then washed twice in cytosolic lysis buffer and pelleted by centrifugation (5,000 g, 10 min, 4 °C) followed by resuspension in nuclear lysis buffer (20 mM HEPES pH 7.9, 40 mM NaCl, 10 % (v/v) glycerol, 50 mM NaF, 5 mM sodium pyrophosphate, 1 mM EDTA, 1.5 mM MgCl_2_, 0.5 mM DTT), mechanically lysed in a Dounce glass homogeniser and incubated on ice for 30 min. The nuclear protein-containing fraction was clarified by centrifugation (16,000 g, 3 min, 4 °C), with all samples flash frozen in L-N_2_ and stored at -80 °C until analysis. All cell lysis buffers were supplemented with a cOmplete protease inhibitor cocktail (Roche).

To purify FLAG-AMPK complexes, they were immobilised by incubation of cell lysates with FLAG-M2 agarose (Sigma-Aldrich A2220) for 2 hours at 4 °C. Following gentle centrifugation, the resin was washed twice in high salt purification buffer (50 mM HEPES pH 7.4, 1 M NaCl, 10 % (v/v) glycerol, 0.02 % (v/v) tween-20) then twice more in the same buffer except with a lower salt concentration (150 mM NaCl).

### Bacterial protein expression and purification

Recombinant full-length heterotrimeric AMPK _6xHis_α11γ1, _6xHis_α12γ1, _6xHis_α21γ1, _6xHis_α2β2γ1 and _6xHis_α2β1γ1 (α2-α1 ST loop chimera) was expressed in *E. coli* Rosetta 2 (DE3) (Merck Millipore) after double-transformation of pET-Duet-1 (α- and γ-subunits) and pCOLA (β-subunit) plasmids and purified as described previously for the _6xHis_α2β1γ1 complex. In brief, expression cultures were grown at 37 °C to an optical density (OD_600_) of 3.0 before induction with 500 μM isopropyl-β-D-1-thiogalactopyranoside (IPTG; Gold Biotechnology) and incubation overnight at 16 °C. Cell pellets were resuspended in lysis buffer (50 mM Tris pH 7.6, 500 mM NaCl, 5 % (v/v) glycerol, 50 mM imidazole, 2 mM β-mercaptoethanol (BME), 0.01 mM leupeptin, 0.1 mM AEBSF, 0.5 mM benzamidine hydrochloride), lysed using a precooled EmulsiFlex-C5 homogeniser (Avestin) and clarified via centrifugation. Protein was bound to a HisTrap HP 5mL Ni^2+^ column (Cytiva) at 1 mL·min^-1^, washed with 10 column volumes of chilled Ni^2+^ column buffer (50 mM Tris pH 7.5, 500 mM NaCl, 5 % (v/v) glycerol, 40 mM imidazole, 2 mM BME) and eluted with Ni^2+^ column buffer supplemented with 400 mM imidazole. Proteinaceous fractions were separated on a HiLoad 16/600 Superdex 200 gel filtration column (Cytiva) pre-equilibrated with AMPK size exclusion column buffer (AMPK SEC buffer; 50 mM Tris pH 8.0, 150 mM NaCl, 2 mM tris(2-carboxyethyl)phosphine (TCEP)). AMPK containing fractions were pooled and concentrated to ∼2 mg·mL^-1^, flash frozen in L-N_2_ and stored at -80 °C. All purified protein was quality controlled by time of flight-mass spectrometry (TOF-MS), SDS-PAGE Coomassie staining and immunoblotting.

### Radioactive kinase assays

AMPK activity was determined as previously described [20]. Briefly, in a 25 μL reaction volume containing 100 μM SAMS peptide (Purar Chemicals, sequence: NH_2_-HMRSAMSGLHLVKRR-COOH), 5 mM MgCl_2_, 200 μM ATP, [γ-^32^P]-ATP (Perkin Elmer), assay buffer (50 mM HEPES pH 7.4, 1 mM DTT and 0.02 % (v/v) tween-20) and purified AMPK, phosphotransferase activity was conducted at 30 °C for 10 minutes and reactions were quenched by spotting 15 μL onto phosphocellulose ion-exchange chromatography paper (prepared in-house). The papers were repeatedly washed in 1 % H_3_PO_4_ (Merck Millipore), added to vials containing 5 mL Ultima Gold liquid scintillation fluid (Perkin Elmer) and the level of ^32^P-transfer to the SAMS peptide was determined using a Tri-Carb 4810TR liquid scintillation counter (Perkin Elmer).

### *In vitro* phosphorylation reactions

Phosphorylation reactions were conducted by incubating bacterially expressed AMPK substrate, 2.5 mM MgCl_2_, 500 μM ATP, AMPK SEC buffer, and mTORC1 complex (Sigma-Aldrich; SRP0364) at a 1:10 kinase:substrate mass ratio. Reactions were quenched by addition of 1 μL of 500 mM EDTA. The quenched reaction mixture was either directly analysed by TOF-MS, digested with trypsin for LC-MS/MS, or diluted with Laemmli sample buffer and 100 ng of substrate analysed by immunoblotting.

### LC-MS analysis on intact protein and peptides

All intact protein and peptide analysis were carried out on a TripleTOF 5600 mass spectrometer (Sciex) operated with the turbo V DuoSpray ion source linked to an Ultimate 3000 RSLCnano system loading pump (Dionex) and Ultimate 3000 RS autosampler (Dionex). The LC-MS was operated using the Analyst TF v1.7.1 software (Sciex). Source and collision gas was provided by a Genius NM3G nitrogen gas generator (PEAK Scientific).

For intact protein analysis (TOF-MS), 5 μg of purified protein was resolved on a Waters Acquity BEH C4 LC guard column (5 mm x 2.1 mm, 1.7 μm, 300 Å). The LC solvent system comprised of H_2_O with 0.1 % formic acid (FA) for channel A, and 90 % acetonitrile/10 % H_2_O with 0.1 % FA for channel B. A flow rate of 50 μL·min^-1^ was used throughout a gradient program consisting of 20 % B (1 minute), 20 to 80 % B (16 minutes), 80 % B (2 minutes), 20 % B (4 minutes). The mass spectrometer was set to TOF-MS acquisition mode and operated in positive ion mode with a mass range of 400-3000 m/z and accumulation time of 0.5 seconds. Spray voltage was set to 5,500 V, source temperature set to 250 °C, ion source gas 1 and 2 set at 35, curtain gas set at 15, and declustering potential set at 180 V. Mass spectra were deconvoluted using Bio Tool Kit in PeakView 2.2 software (Sciex).

To prepare FLAG immobilised AMPK for peptide analysis, protein was precipitated by addition of 1 mL 100 % methanol to the resin (10x resin volume) and incubation on ice for 30 mins. The sample was centrifuged (12,000 g, 10 min, 4 °C), supernatant discarded, and resin dried under N_2_ gas. The dried resin was resuspended with 50 mM Tris pH 7.5 (2x resin volume) and 200 ng of trypsin was added (Promega; made up to 100 ng·μL^-1^ in H_2_O), the samples were digested overnight shaking at 37°C. To prepare purified AMPK (*E. coli* preps) for peptide analysis, 20 μg of protein was diluted in 50 mM Tris pH 7.5 and 200 ng of Trypsin was added, the samples were digested overnight at 37 °C. All digests were quenched by addition of 1 μL of 100 % FA before supernatant was removed for LC-MS/MS analysis.

Digested AMPK was resolved on a Waters Acquity BEH peptide C18 column (100 mm x 2.1 mm, 1.7 μm, 130 Å). The LC solvent system comprised of H_2_O with 0.1 % FA for channel A, and 90 % acetonitrile/10 % H_2_O with 0.1 % FA for channel B. A flow rate of 125 μL·min^-1^ was used throughout a gradient program consisting of 2.5 % B (2.5 minutes), 2.5 to 60 % B (67.5 minutes), 60 to 100 % B (20 minutes), 100 % B (10 minutes), 100 to 2.5 % B (2 minutes), 2.5 % B (8 minutes). The mass spectrometer was set to time of flight-mass spectrometry (TOF-MS) acquisition mode and operated in positive ion mode with a mass range of 200-3000 m/z and accumulation time of 0.5 seconds. Spray voltage was set to 5,500 V, source temperature set to 300 °C, ion source gas 1 and 2 set at 35, curtain gas set at 15, and declustering potential set at 150 V. TOF-MS data was visualized using PeakView v2.2 software (Sciex) and peptide peak areas quantified using Skyline v21.1 (MacCross Lab Software). Phosphorylation stoichiometries were calculated from phosphorylated and dephosphorylated peptide peak areas using Equation 1.

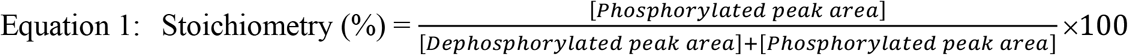

### Cell proliferation assays

Cell proliferation assays were conducted as described previously [27]. Briefly, 24 hours after HEK293 cells were transfected with heterotrimeric AMPK (GFP-α1/2, β1-myc, HA-γ1; wild-type and indicated mutants), the media was replaced with either fresh DMEM (supplemented with 10 % FBS, 4 mM L-glutamine and penicillin-streptomycin) or arginine- and lysine-free DMEM (supplemented with dialysed 10 % FBS and penicillin-streptomycin). Cell proliferation was tracked in real-time using the Incucyte® Live-Cell Analysis System (Sartorius) according to the manufacturer’s instructions.

### Statistical analysis

All statistical analyses were performed using Prism v9.2.0 (GraphPad Software). Results from replicate experiments (n) are expressed as means +/-standard error (SEM).

## References

1 Wang, Y.-P. and Lei, Q.-Y. (2018) Metabolite sensing and signaling in cell metabolism. Signal transduction and targeted therapy, Springer US 3, 30.

2 Milanesi, R., Coccetti, P. and Tripodi, F. (2020) The Regulatory Role of Key Metabolites in the Control of Cell Signaling. Biomolecules 10, 862.

3 Kwon, Y. T. and Ciechanover, A. (2017) The Ubiquitin Code in the Ubiquitin-Proteasome System and Autophagy. Trends in Biochemical Sciences, Elsevier Ltd 42, 873–886.

4 Hardie, D. G. and Carling, D. (1997) The AMP-activated protein kinase. Fuel gauge of the mammalian cell? European Journal of Biochemistry 246, 259–273.

5 Carling, D. (2004) The AMP-activated protein kinase cascade – a unifying system for energy control. Trends in Biochemical Sciences 29, 18–24.

6 Oakhill, J. S., Scott, J. W. and Kemp, B. E. (2009) Structure and function of AMP-activated protein kinase. Acta Physiologica 196, 3–14.

7 Uhlen, M., Fagerberg, L., Hallstrom, B. M., Lindskog, C., Oksvold, P., Mardinoglu, A., Sivertsson, A., Kampf, C., Sjostedt, E., Asplund, A., et al. (2015) Tissue-based map of the human proteome. Science 347, 1260419–1260419.

8 Hawley, S. A., Davison, M., Woods, A., Davies, S. P., Beri, R. K., Carling, D. and Hardie, D. G. (1996) Characterization of the AMP-activated Protein Kinase Kinase from Rat Liver and Identification of Threonine 172 as the Major Site at Which It Phosphorylates AMP-activated Protein Kinase. Journal of Biological Chemistry 271, 27879–27887.

9 Hawley, S. A., Boudeau, J., Reid, J. L., Mustard, K. J., Udd, L., Mäkelä, T. P., Alessi, D. R. and Hardie, D. G. (2003) Complexes between the LKB1 tumor suppressor, STRAD alpha/beta and MO25 alpha/beta are upstream kinases in the AMP-activated protein kinase cascade. Journal of biology 2, 28.

10 Shaw, R. J., Kosmatka, M., Bardeesy, N., Hurley, R. L., Witters, L. A., DePinho, R. A. and Cantley, L. C. (2004) The tumor suppressor LKB1 kinase directly activates AMP-activated kinase and regulates apoptosis in response to energy stress. Proceedings of the National Academy of Sciences 101, 3329–3335.

11 Hurley, R. L., Anderson, K. A., Franzone, J. M., Kemp, B. E., Means, A. R. and Witters, L.A. (2005) The Ca 2+ /Calmodulin-dependent Protein Kinase Kinases Are AMP-activated Protein Kinase Kinases. Journal of Biological Chemistry 280, 29060–29066.

12 Ovens, A. J., Scott, J. W., Langendorf, C. G., Kemp, B. E., Oakhill, J. S. and Smiles, W. J. (2021) Post-Translational Modifications of the Energy Guardian AMP-Activated Protein Kinase. International journal of molecular sciences 22, 1229.

13 Yan, Y., Zhou, X. E., Xu, H. E. and Melcher, K. (2018) Structure and physiological regulation of AMPK. International Journal of Molecular Sciences 19.

14 Ngoei, K. R. W., Langendorf, C. G., Ling, N. X. Y., Hoque, A., Varghese, S., Camerino, M. A., Walker, S. R., Bozikis, Y. E., Dite, T. A., Ovens, A. J., et al. (2018) Structural Determinants for Small-Molecule Activation of Skeletal Muscle AMPK α2β2γ1 by the Glucose Importagog SC4. Cell chemical biology 25, 728-737.e9.

15 Myers, R. W., Guan, H.-P., Ehrhart, J., Petrov, A., Prahalada, S., Tozzo, E., Yang, X., Kurtz, M. M., Trujillo, M., Gonzalez Trotter, D., et al. (2017) Systemic pan-AMPK activator MK-8722 improves glucose homeostasis but induces cardiac hypertrophy. Science 357, 507–511.

16 Cokorinos, E. C., Delmore, J., Reyes, A. R., Albuquerque, B., Kjøbsted, R., Jørgensen, N. O., Tran, J.-L., Jatkar, A., Cialdea, K., Esquejo, R. M., et al. (2017) Activation of Skeletal Muscle AMPK Promotes Glucose Disposal and Glucose Lowering in Non-human Primates and Mice. Cell metabolism, Elsevier Inc. 25, 1147-1159.e10.

17 Pinkosky, S. L., Scott, J. W., Desjardins, E. M., Smith, B. K., Day, E. A., Ford, R. J., Langendorf, C. G., Ling, N. X. Y., Nero, T. L., Loh, K., et al. (2020) Long-chain fatty acyl-CoA esters regulate metabolism via allosteric control of AMPK β1 isoforms. Nature Metabolism, Springer US.

18 Davies, S. P., Helps, N. R., Cohen, P. T. W. and Hardie, D. G. (1995) 5’-AMP inhibits dephosphorylation, as well as promoting phosphorylation, of the AMP-activated protein kinase. Studies using bacterially expressed human protein phosphatase-2C alpha and native bovine protein phosphatase-2AC. FEBS letters 377, 421–5.

19 Ross, F. A., Jensen, T. E. and Hardie, D. G. (2016) Differential regulation by AMP and ADP of AMPK complexes containing different γ subunit isoforms. Biochemical Journal 473, 189– 199.

20 Scott, J. W., Ling, N., Issa, S. M. A., Dite, T. A., O’Brien, M. T., Chen, Z.-P., Galic, S., Langendorf, C. G., Steinberg, G. R., Kemp, B. E., et al. (2014) Small molecule drug A-769662 and AMP synergistically activate naive AMPK independent of upstream kinase signaling. Chemistry & biology, Elsevier Ltd 21, 619–27.

21 Rogala, K. B., Gu, X., Kedir, J. F., Abu-Remaileh, M., Bianchi, L. F., Bottino, A. M. S., Dueholm, R., Niehaus, A., Overwijn, D., Fils, A.-C. P., et al. (2019) Structural basis for the docking of mTORC1 on the lysosomal surface. Science (New York, N.Y.) 366, 468–475.

22 Sancak, Y., Peterson, T. R., Shaul, Y. D., Lindquist, R. A., Thoreen, C. C., Bar-Peled, L. and Sabatini, D. M. (2008) The Rag GTPases bind raptor and mediate amino acid signaling to mTORC1. Science (New York, N.Y.) 320, 1496–501.

23 Sancak, Y., Bar-Peled, L., Zoncu, R., Markhard, A. L., Nada, S. and Sabatini, D. M. (2010) Ragulator-Rag complex targets mTORC1 to the lysosomal surface and is necessary for its activation by amino acids. Cell, Elsevier B.V. 141, 290–303.

24 Menon, S., Dibble, C. C., Talbott, G., Hoxhaj, G., Valvezan, A. J., Takahashi, H., Cantley, L. C. and Manning, B. D. (2014) Spatial control of the TSC complex integrates insulin and nutrient regulation of mTORC1 at the lysosome. Cell, Elsevier B.V. 156, 771–85.

25 Gwinn, D. M., Shackelford, D. B., Egan, D. F., Mihaylova, M. M., Mery, A., Vasquez, D. S., Turk, B. E. and Shaw, R. J. (2008) AMPK Phosphorylation of Raptor Mediates a Metabolic Checkpoint. Molecular Cell 30, 214–226.

26 Inoki, K., Zhu, T. and Guan, K.-L. (2003) TSC2 Mediates Cellular Energy Response to Control Cell Growth and Survival. Cell 115, 577–590.

27 Ling, N. X. Y., Kaczmarek, A., Hoque, A., Davie, E., Ngoei, K. R. W., Morrison, K. R., Smiles, W. J., Forte, G. M., Wang, T., Lie, S., et al. (2020) mTORC1 directly inhibits AMPK to promote cell proliferation under nutrient stress. Nature metabolism, Nature Research 2, 41–49.

28 Dickhut, C., Feldmann, I., Lambert, J. and Zahedi, R. P. (2014) Impact of Digestion Conditions on Phosphoproteomics. Journal of Proteome Research 13, 2761–2770.

29 Bubis, J. A., Gorshkov, V., Gorshkov, M. v. and Kjeldsen, F. (2020) PhosphoShield: Improving Trypsin Digestion of Phosphoproteins by Shielding the Negatively Charged Phosphate Moiety. Journal of the American Society for Mass Spectrometry 31, 2053–2060.

30 Rajamohan, F., Reyes, A. R., Frisbie, R. K., Hoth, L. R., Sahasrabudhe, P., Magyar, R., Landro, J. A., Withka, J. M., Caspers, N. L., Calabrese, M. F., et al. (2016) Probing the enzyme kinetics, allosteric modulation and activation of α1- and α2-subunit-containing AMP-activated protein kinase (AMPK) heterotrimeric complexes by pharmacological and physiological activators. The Biochemical journal 473, 581–92.

31 Willows, R., Navaratnam, N., Lima, A., Read, J. and Carling, D. (2017) Effect of different γ-subunit isoforms on the regulation of AMPK. Biochemical Journal 474, 1741–1754.

32 Mitchelhill, K. I., Michell, B. J., House, C. M., Stapleton, D., Dyck, J., Gamble, J., Ullrich, C., Witters, L. A. and Kemp, B. E. (1997) Posttranslational Modifications of the 5′-AMP-activated Protein Kinase β1 Subunit. Journal of Biological Chemistry 272, 24475–24479.

33 Dite, T. A., Ling, N. X. Y., Scott, J. W., Hoque, A., Galic, S., Parker, B. L., Ngoei, K. R. W., Langendorf, C. G., O’Brien, M. T., Kundu, M., et al. (2017) The autophagy initiator ULK1 sensitizes AMPK to allosteric drugs. Nature communications, Springer US 8, 571.

34 Horman, S., Vertommen, D., Heath, R., Neumann, D., Mouton, V., Woods, A., Schlattner, U., Wallimann, T., Carling, D., Hue, L., et al. (2006) Insulin Antagonizes Ischemia-induced Thr 172 Phosphorylation of AMP-activated Protein Kinase α-Subunits in Heart via Hierarchical Phosphorylation of Ser 485/491. Journal of Biological Chemistry 281, 5335– 5340.

35 Hawley, S. A., Ross, F. A., Gowans, G. J., Tibarewal, P., Leslie, N. R. and Hardie, D. G. (2014) Phosphorylation by Akt within the ST loop of AMPK-α1 down-regulates its activation in tumour cells. The Biochemical journal 459, 275–87.

36 Suzuki, T., Bridges, D., Nakada, D., Skiniotis, G., Morrison, S. J., Lin, J. D., Saltiel, A. R. and Inoki, K. (2013) Inhibition of AMPK catabolic action by GSK3. Molecular cell, Elsevier Inc. 50, 407–19.

37 Djouder, N., Tuerk, R. D., Suter, M., Salvioni, P., Thali, R. F., Scholz, R., Vaahtomeri, K., Auchli, Y., Rechsteiner, H., Brunisholz, R. A., et al. (2010) PKA phosphorylates and inactivates AMPKα to promote efficient lipolysis. EMBO Journal 29, 469–481.

38 Kuenzel, E. A., Mulligan, J. A., Sommercorn, J. and Krebs, E. G. (1987) Substrate specificity determinants for casein kinase II as deduced from studies with synthetic peptides. The Journal of biological chemistry 262, 9136–40.

39 Kemp, B. E., Graves, D. J., Benjamini, E. and Krebs, E. G. (1977) Role of multiple basic residues in determining the substrate specificity of cyclic AMP-dependent protein kinase. The Journal of biological chemistry 252, 4888–94.

40 Hurley, R. L., Barré, L. K., Wood, S. D., Anderson, K. A., Kemp, B. E., Means, A. R. and Witters, L. A. (2006) Regulation of AMP-activated protein kinase by multisite phosphorylation in response to agents that elevate cellular cAMP. The Journal of biological chemistry 281, 36662–72.

41 Heathcote, H. R., Mancini, S. J., Strembitska, A., Jamal, K., Reihill, J. A., Palmer, T. M., Gould, G. W. and Salt, I. P. (2016) Protein kinase C phosphorylates AMP-activated protein kinase α1 Ser487. Biochemical Journal 473, 4681–4697.

42 Dagon, Y., Hur, E., Zheng, B., Wellenstein, K., Cantley, L. C. and Kahn, B. B. (2012) p70S6 Kinase Phosphorylates AMPK on Serine 491 to Mediate Leptin’s Effect on Food Intake. Cell Metabolism, Elsevier Inc. 16, 104–112.

43 Coughlan, K. A., Valentine, R. J., Sudit, B. S., Allen, K., Dagon, Y., Kahn, B. B., Ruderman, N. B. and Saha, A. K. (2016) PKD1 Inhibits AMPKα2 through Phosphorylation of Serine 491 and Impairs Insulin Signaling in Skeletal Muscle Cells. Journal of Biological Chemistry 291, 5664–5675.

44 Stauffer, S., Zeng, Y., Santos, M., Zhou, J., Chen, Y. and Dong, J. (2019) Cyclin-dependent kinase 1-mediated AMPK phosphorylation regulates chromosome alignment and mitotic progression. Journal of Cell Science, Company of Biologists Ltd 132.

45 Warden, S. M., Richardson, C., O’Donnell J.J., Stapleton, D. I., Kemp, B. E. and Witters, L.A. (2001) Post-translational modifications of the β-1 subunit of AMP-activated protein kinase affect enzyme activity and cellular localization. Biochemical Journal 354, 275–283.

46 Chen, Z., Heierhorst, J., Mann, R. J., Mitchelhill, K. I., Michell, B. J., Witters, L. A., Lynch, G. S., Kemp, B. E. and Stapleton, D. (1999) Expression of the AMP-activated protein kinase β1 and β2 subunits in skeletal muscle. FEBS Letters 460, 343–348.

47 Hornbeck, P. v., Kornhauser, J. M., Tkachev, S., Zhang, B., Skrzypek, E., Murray, B., Latham, V. and Sullivan, M. (2012) PhosphoSitePlus: a comprehensive resource for investigating the structure and function of experimentally determined post-translational modifications in man and mouse. Nucleic Acids Research 40, D261–D270.

48 Brunn, G. J., Fadden, P., Haystead, T. A. and Lawrence, J. C. (1997) The mammalian target of rapamycin phosphorylates sites having a (Ser/Thr)-Pro motif and is activated by antibodies to a region near its COOH terminus. The Journal of biological chemistry 272, 32547–50.

49 Rajamohan, F., Harris, M. S., Frisbie, R. K., Hoth, L. R., Geoghegan, K. F., Valentine, J. J., Reyes, A. R., Landro, J. A., Qiu, X. and Kurumbail, R. G. (2010) Escherichia coli expression, purification and characterization of functional full-length recombinant α2β2γ3 heterotrimeric complex of human AMP-activated protein kinase. Protein Expression and Purification, Elsevier Inc. 73, 189–197.

50 Nelson, M. E., Parker, B. L., Burchfield, J. G., Hoffman, N. J., Needham, E. J., Cooke, K. C., Naim, T., Sylow, L., Ling, N. X., Francis, D., et al. (2019) Phosphoproteomics reveals conserved exercise-stimulated signaling and AMPK regulation of store-operated calcium entry. The EMBO journal 38, e102578.

51 Thoreen, C. C., Kang, S. A., Chang, J. W., Liu, Q., Zhang, J., Gao, Y., Reichling, L. J., Sim, T., Sabatini, D. M. and Gray, N. S. (2009) An ATP-competitive mammalian target of rapamycin inhibitor reveals rapamycin-resistant functions of mTORC1. The Journal of biological chemistry, American Society for Biochemistry and Molecular Biology Inc. 284, 8023–32.

52 Facchinetti, V., Ouyang, W., Wei, H., Soto, N., Lazorchak, A., Gould, C., Lowry, C., Newton, A. C., Mao, Y., Miao, R. Q., et al. (2008) The mammalian target of rapamycin complex 2 controls folding and stability of Akt and protein kinase C. The EMBO journal 27, 1932–43.

53 Oh, W. J., Wu, C., Kim, S. J., Facchinetti, V., Julien, L.-A., Finlan, M., Roux, P. P., Su, B. and Jacinto, E. (2010) mTORC2 can associate with ribosomes to promote cotranslational phosphorylation and stability of nascent Akt polypeptide. The EMBO journal 29, 3939–51.

54 Suzuki, A., Okamoto, S., Lee, S., Saito, K., Shiuchi, T. and Minokoshi, Y. (2007) Leptin Stimulates Fatty Acid Oxidation and Peroxisome Proliferator-Activated Receptor α Gene Expression in Mouse C2C12 Myoblasts by Changing the Subcellular Localization of the α2 Form of AMP-Activated Protein Kinase. Molecular and Cellular Biology 27, 4317–4327.

55 Lamia, K. A., Sachdeva, U. M., DiTacchio, L., Williams, E. C., Alvarez, J. G., Egan, D. F., Vasquez, D. S., Juguilon, H., Panda, S., Shaw, R. J., et al. (2009) AMPK regulates the circadian clock by cryptochrome phosphorylation and degradation. Science (New York, N.Y.) 326, 437–40.

56 Inoki, K., Li, Y., Zhu, T., Wu, J. and Guan, K.-L. (2002) TSC2 is phosphorylated and inhibited by Akt and suppresses mTOR signalling. Nature cell biology 4, 648–57.

57 Vara-Ciruelos, D., Russell, F. M. and Hardie, D. G. (2019) The strange case of AMPK and cancer: Dr Jekyll or Mr Hyde? †. Open biology 9, 190099.

58 Vega-Rubin-de-Celis, S., Peña-Llopis, S., Konda, M. and Brugarolas, J. (2017) Multistep regulation of TFEB by MTORC1. Autophagy, Taylor and Francis Inc. 13, 464–472.

59 Napolitano, G., Esposito, A., Choi, H., Matarese, M., Benedetti, V., di Malta, C., Monfregola, J., Medina, D. L., Lippincott-Schwartz, J. and Ballabio, A. (2018) mTOR-dependent phosphorylation controls TFEB nuclear export. Nature communications, Nature Publishing Group 9, 3312.

60 Settembre, C., Zoncu, R., Medina, D. L., Vetrini, F., Erdin, S., Erdin, S., Huynh, T., Ferron, M., Karsenty, G., Vellard, M. C., et al. (2012) A lysosome-to-nucleus signalling mechanism senses and regulates the lysosome via mTOR and TFEB. The EMBO journal 31, 1095–108.

61 Napolitano, G., di Malta, C., Esposito, A., de Araujo, M. E. G., Pece, S., Bertalot, G., Matarese, M., Benedetti, V., Zampelli, A., Stasyk, T., et al. (2020) A substrate-specific mTORC1 pathway underlies Birt-Hogg-Dubé syndrome. Nature, Nature Research 585, 597– 602.

62 Paquette, M., El-Houjeiri, L.C, Zirden, L., Puustinen, P., Blanchette, P., Jeong, H., Dejgaard, K., Siegel, P. M. and Pause, A. (2021) AMPK-dependent phosphorylation is required for transcriptional activation of TFEB and TFE3. Autophagy, Springer US 11, 1–19.

63 Zhou, X., Li, S. and Zhang, J. (2016) Tracking the Activity of mTORC1 in Living Cells Using Genetically Encoded FRET-based Biosensor TORCAR. Current protocols in chemical biology 8, 225–233.

64 Zhou, X., Zhong, Y., Molinar-Inglis, O., Kunkel, M. T., Chen, M., Sun, T., Zhang, J., Shyy, J. Y.-J., Trejo, J., Newton, A. C., et al. (2020) Location-specific inhibition of Akt reveals regulation of mTORC1 activity in the nucleus. Nature communications 11, 6088.

65 Blair, E., Redwood, C., Ashrafian, H., Oliveira, M., Broxholme, J., Kerr, B., Salmon, A., Ostman-Smith, I. and Watkins, H. (2001) Mutations in the gamma(2) subunit of AMP-activated protein kinase cause familial hypertrophic cardiomyopathy: evidence for the central role of energy compromise in disease pathogenesis. Human molecular genetics 10, 1215–20.

66 Gollob, M. H., Green, M. S., Tang, A. S., Gollob, T., Karibe, A., Ali Hassan, A. S., Ahmad, F., Lozado, R., Shah, G., Fananapazir, L., et al. (2001) Identification of a gene responsible for familial Wolff-Parkinson-White syndrome. The New England journal of medicine 344, 1823–31.

67 Yavari, A., Bellahcene, M., Bucchi, A., Sirenko, S., Pinter, K., Herring, N., Jung, J. J., Tarasov, K. v, Sharpe, E. J., Wolfien, M., et al. (2017) Mammalian γ2 AMPK regulates intrinsic heart rate. Nature communications 8, 1258.

68 Scott, J. W., van Denderen, B. J. W., Jorgensen, S. B., Honeyman, J. E., Steinberg, G. R., Oakhill, J. S., Iseli, T. J., Koay, A., Gooley, P. R., Stapleton, D., et al. (2008) Thienopyridone Drugs Are Selective Activators of AMP-Activated Protein Kinase β1-Containing Complexes. Chemistry and Biology, Elsevier Ltd 15, 1220–1230.

69 Oakhill, J. S., Scott, J. W. and Dite, T. A. (2018) Transient Expression of AMPK Heterotrimer Complexes in Mammalian Cells. In Methods in Molecular Biology, pp 159– 169.

